# The inflammasome of circulatory collapse: single cell analysis of survival on extracorporeal life support

**DOI:** 10.1101/568659

**Authors:** Eric J. Kort, Matthew Weiland, Edgars Grins, Emily Eugster, Hsiao-yun Milliron, Catherine Kelty, Nabin Manandhar Shrestha, Tomasz Timek, Marzia Leacche, Stephen J Fitch, Theodore J Boeve, Greg Marco, Michael Dickinson, Penny Wilton, Stefan Jovinge

## Abstract

**Background:** Despite being a lifesaving intervention for the most critically ill and circulatory compromised patients, veno-arterial extra-corporeal life support (VA-ECLS) is associated with a mortality rate of nearly 60%. Understanding how the immune response to VA-ECLS either promotes or impedes survival would both enhance risk stratification and uncover new therapeutic strategies for these patients. However, conventional enumeration of peripheral blood mononuclear cells (PBMCs) and their subsets have failed to identify determinants of outcome among these cells.

**Methods:** Flow cytometry and plasma cytokine measurement was combined with single cell RNASeq analysis of PBMCs from patients in circulatory shock being started on VA-ECLS to identify clinical, laboratory, and cellular features associated with 72 hour survival.

**Results:** Non-surviving patients exhibited higher plasma levels of the tissue aggressive inflammatory cytokines IL-1, IL-6, IL-12 and TNF-α. Distribution of cells between conventional PBMC subtypes was not predictive of survival. Single cell RNASeq analysis of discriminatory markers within each PBMC subtype revealed that the proportion of CD8^+^ Natural Killer T-cells (NKT) that expressed CD52, a known immune-modulator, was associated with improved survival. This cell population correlated inversely with IL-6 production. CD8^+^/CD52^+^ NKT cells were quantified by flow cytometry in a second, validation cohort. Those patients with a high proportion of CD52+ cells among all CD8^+^ NKT cells had more severe disease relative to the low CD52+ group, but nevertheless were nearly 5 time less likely to die in the first 72 hours of VA-ECLS (p=0.043 by log rank test).

**Conclusions:** CD8^+^/CD52^+^ NKT cells are associated with survival in patients undergoing VA-ECLS. Fluidics based scRNASeq can reveal important aspects of pathophysiology in complex disease states such as circulatory collapse and VA-ECLS. Further studies in animal models will be required to determine if stimulation of CD8^+^/CD52^+^ NKT cell expansion may be an effective therapeutic strategy in this patient population.

## Introduction

Cardiac injury may culminate in circulatory collapse requiring prompt intervention including veno-arterial extra-corporeal life support (VA-ECLS). VA-ECLS, also referred to as veno-arterial extra corporeal membrane oxygenation, represents a lifesaving approach to lung and/or heart bypass for critically ill patients. However, patients requiring VA-ECLS have a survival to discharge rate of only about 40% (*1*) - a fact that, while sobering, is not surprising given the dire clinical condition of patients requiring VA-ECLS. Mortality while on VA-ECLS is due not only to the relentless progression of the underlying disease processes, but also complications of VA-ECLS itself. These complications include derangement of clotting and the inflammatory stimulation produced by the foreign material surfaces and mechanical stresses presented by the bypass circuit. Understanding how the immune response in the context of circulatory collapse and VA-ECLS either promotes or impedes survival would both enhance risk stratification and uncover new therapeutic strategies for these patients. Here we characterized the inflammatory milieu of circulatory collapse in terms of cytokine release to identify factors associated with survival among these cell-signaling molecules. We then used single cell transcriptomics in an effort to identify what cell populations may be key modulators of this inflammatory response as it relates to survival.

Conventional enumeration of peripheral blood mononuclear cells (PBMCs) and their subsets have failed to identify determinants of outcome among these cells for this complex patient population. Single cell transcriptomic approaches continue to modify and expand our understanding of the function and classification of PBMC subsets (*2, 3*). The degree to which these sophisticated approaches will lead to clinically actionable information remains to be established (*4*). To that end, we here took an integrated approach combining plasma cytokine quantification, flow cytometry, and single cell transcriptomics to address the challenging question of why some patients survive on VA-ECLS while many do not. We profiled approximately 40,000 cells from 38 patients who had undergone VA-ECLS due to decompensated heart failure. First, we assigned cells to their canonical PBMC types based on RNA expression levels of established surface markers. We then validated these assignments using conventional flow cytometry and interrogated each of these subpopulations for survival markers across the genome. Finally, we validated our findings in a separate cohort of patients by flow cytometry.

## Results

### Clinical characteristics of cohort

For this study, clinical data and blood samples were obtained prospectively from 38 patients being started on VA-ECLS due to decompensated heart failure. We analyzed clinical and laboratory parameters that were predictive of survival through the first 72 hours of VA-ECLS in these patients. Consistent with prior literature (*5*), non-surviving patients exhibited more acidosis, higher SOFA score, and worse renal function relative to surviving patients (**Table 1** & **Fig 1A**). Age was not predictive of survival in this cohort.

**Table 1:** Clinical characteristics of study participants. “t” = t-test, Wilc=Wilcox test.

| Variable | Survived |  |  | Died |  |  | p-value | Test |
| --- | --- | --- | --- | --- | --- | --- | --- | --- |
| | N | Mean ± SD | Min - Max | N | Mean ± SD | Min - Max | | |
| Age (yr) | 26 | 60.42 ± 13.33 | 24 - 82 | 12 | 58.08 ± 13.56 | 34 - 77 | 0.62 | t |
| Height (cm) | 26 | 173.4 ± 7.75 | 155 - 190.25 | 12 | 175.74 ± 13.53 | 147.32 - 193.02 | 0.502 | t |
| Weight (kg) | 26 | 88.66 ± 12.89 | 70 - 123 | 12 | 101.32 ± 22.84 | 60.8 - 141 | 0.094 | t |
| BMI (kg/m<sup>2</sup>) | 26 | 29.58 ± 4.5 | 20.9 - 40.2 | 12 | 32.57 ± 5.48 | 26.7 - 44.5 | 0.084 | t |
| Albumin (g/dL) | 26 | 2.66 ± 0.51 | 1.4 - 3.6 | 12 | 2.05 ± 0.45 | 1.3 - 2.7 | 0.001 | t |
| AST (IU/L) | 26 | 369.62 ± 860.21 | 20 - 4325 | 12 | 1554.17 ± 2200 | 23 - 6858 | 0.14 | Wilc |
| ALT (IU/L) | 26 | 308.15 ± 981.43 | 7-4997 | 12 | 707.5 ± 995.32 | 16 - 2945 | 0.106 | Wilc |
| Bilirubin Total (mg/dL) | 26 | 1.2 ± 0.84 | 0.3 - 3.9 | 12 | 1.89 ± 2.83 | 0.2 - 10.7 | 0.705 | Wilc |
| pH Arterial | 26 | 7.34 ± 0.13 | 7.1 - 7.6 | 12 | 7.15 ± 0.14 | 6.91 - 7.35 | <0.001 | t |
| Lactate (mmol/L) | 26 | 5.62 ± 3.49 | 1 - 13.5 | 12 | 11.84 ± 4.86 | 2.3 - 19.8 | <0.001 | t |
| Creatinine (mg/dL) | 26 | 1.24 ± 0.36 | 0.61 - 2.42 | 12 | 1.68 ± 0.58 | 1.1 - 2.91 | 0.031 | Wilc |
| MDRD eGFR (ml/min/1.73 m<sup>2</sup>) | 26 | 63.88 ± 23.38 | 27 - 130.68 | 12 | 42.91 ± 14.36 | 23 - 64.3 | 0.007 | t |
| Sodium (mmol/L) | 26 | 141.96 ± 5.74 | 122 - 153 | 12 | 145.75 ± 7.56 | 137 - 162 | 0.344 | Wilc |
| Potassium (mmol/L) | 26 | 3.75 ± 0.56 | 2.6 - 5 | 12 | 4.28 ± 0.74 | 3.2 - 5.9 | 0.021 | t |
| Magnesium (mg/dL) | 26 | 2.13 ± 0.4 | 1.5 - 3.2 | 11 | 2.43 ± 0.68 | 1.7 - 3.6 | 0.198 | t |
| Glucose (mg/dL) | 26 | 203.19 ± 81.08 | 82 - 385 | 12 | 212.25 ± 95.21 | 97 - 410 | 0.764 | t |
| CVP(RAP) (mmHg) | 26 | 12.58 ± 6.28 | 0 - 25 | 11 | 13.36 ± 6.28 | 2-21 | 0.73 | t |
| MAP Arterial (mmHg) | 26 | 78.54 ± 14.98 | 50 - 112 | 12 | 63.17 ± 12.07 | 30 - 76 | 0.004 | Wilc |
| SVR | 17 | 1317.06 ± 901.84 | 568 - 3509 | 2 | 1036 ± 1047.93 | 295 - 1777 | 0.77 | t |
| SBP Arterial (mmHg) | 26 | 101.31 ± 25.59 | 65 - 167 | 11 | 85.64 ± 24.05 | 32 - 116 | 0.189 | Wilc |
| DBP Arterial (mmHg) | 26 | 71.81 ± 21.89 | 42 - 154 | 11 | 61.27 ± 19.66 | 29 - 107 | 0.122 | Wilc |
| Core Temperature (0C) | 26 | 36.14 ± 0.91 | 33.8 - 37.6 | 10 | 35.94 ± 1.04 | 34.5 - 37.4 | 0.671 | Wilc |
| Inotrope Score | 26 | 18.44 ± 12.49 | 0 - 55 | 12 | 32.9 ± 30.68 | 4 - 120 | 0.087 | Wilc |
| SOFA score | 26 | 10.08 ± 1.52 | 7-13 | 11 | 11.82 ± 2.48 | 8-16 | 0.013 | t |

**Figure 1:**
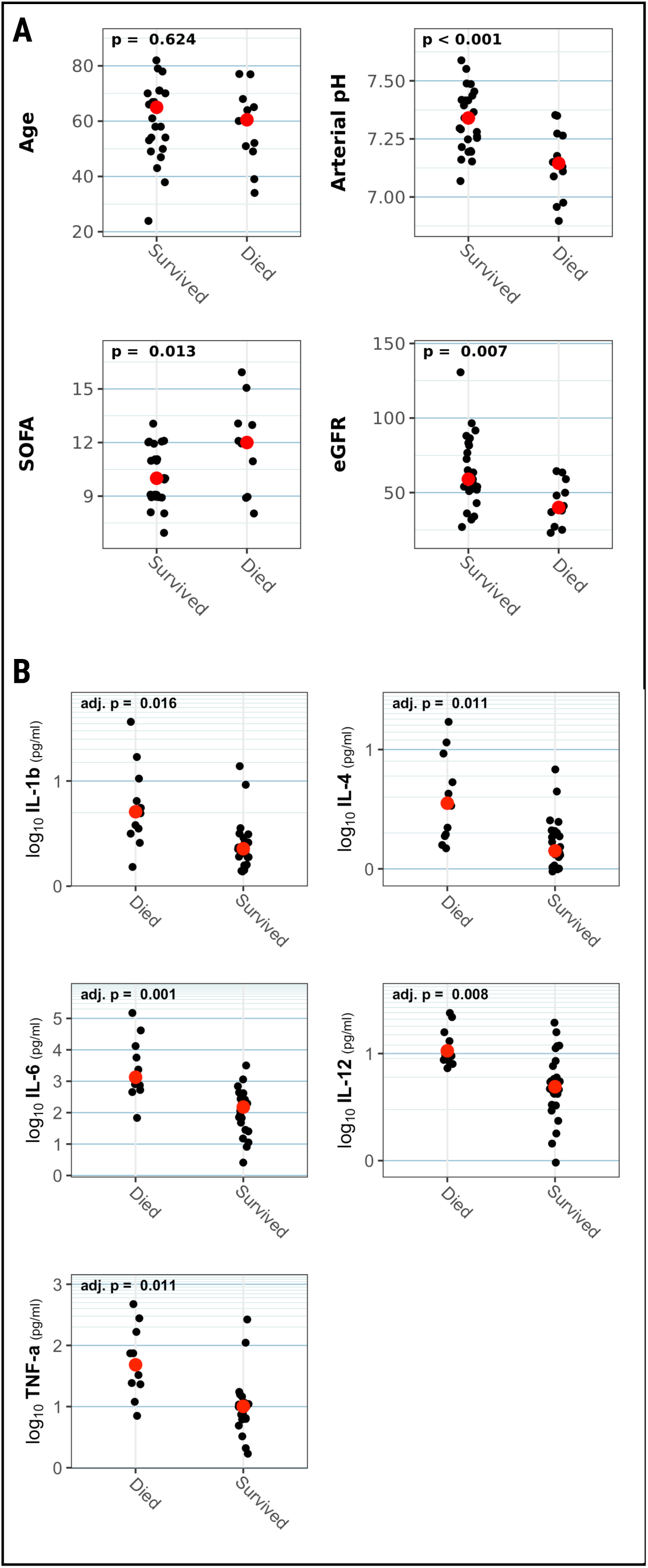
Clinical and laboratory characteristics of surviving and non-surviving patients. **(A)** Key clinical parameters of patients, stratified by 72 hour survival (N=38, p-values are for t-tests comparing survived and non-survived). **(B)** Plasma cytokine levels exhibiting significant differences between surviving and non-surviving (72 hour) patients (N=36, p-values are for Wilcoxon rank sum test adjusted for multiple comparisons by method of Holm).

### Inflammatory cytokines predictive of mortality

We quantified the serum levels of 17 cytokines in 38 patients requiring VA-ECLS due to decompensated heart failure (supplemental **Table S1**). After adjustment for multiple comparisons, 5 of the 17 were significantly associated with survival (**Fig. 1B**). Specifically, the cytokines IL-1β, IL-4, IL-6, IL-12, and TNAα were all higher in patients who did not survive vs. those who survived the first 72 hours of VA-ECLS.

While it is noteworthy that pro-inflammatory cytokines are a marker of poor outcome in these patients, we wanted to gain additional insight into what components of the immune response might be driving these and related components of the inflammatory response. In order to determine what cell populations might be responsible for the differential inflammatory response between surviving and non-surviving patients, we performed scRNASeq based transcriptional profiling of a total of 40,000 peripheral blood mononuclear cells from these patients (mean time between sample acquisition and VA-ECLS, ±79 minutes) on the inDrop microfluidics encapsulation platform (*6*). We were interested in identifying whether scRNASeq analysis combined with both cytokine levels and flow cytometric data (**Fig. 2A**) could provide additional predictors of—and potentially mechanistic insights into—survival in these patients.

**Figure 2.**
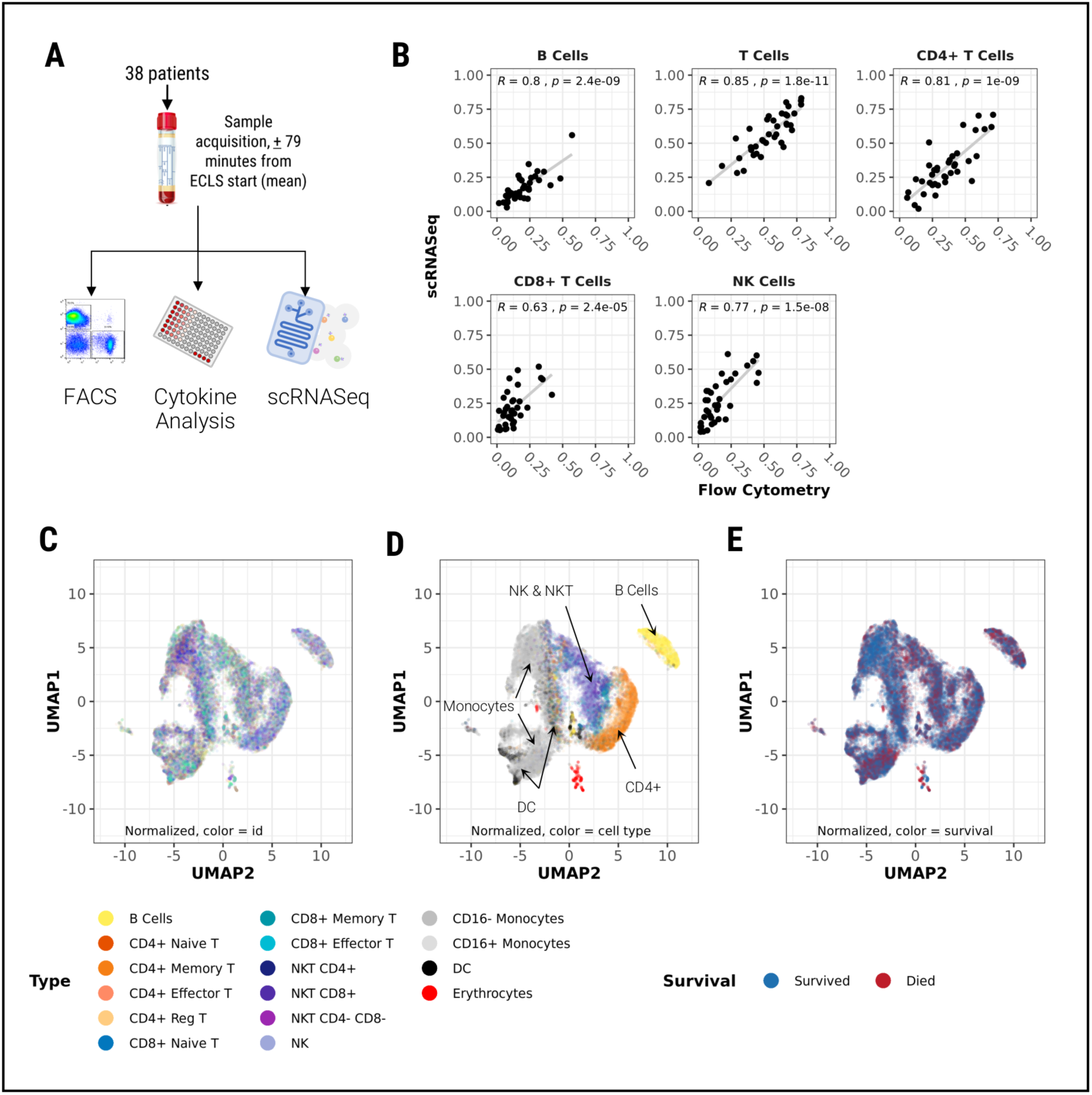
Single cell analysis of PBMCs at time of VA-ECLS initiation. **(A)** Overview of study design. **(B)** Validation of cell type assignment by RNA expression of canonical surface markers. Major lymphocyte populations were quantified by both conventional flow cytometry and scRNASeq analysis. The proportion of cells in each population (as a proportion of all lymphocytes) was compared for each patient between the two modalities. Assignments showed good correlation between conventional flow cytometry and scRNASeq (N=38, p-value is for Pearson’s correlation coefficient). **(C)** Unsupervised clustering by UMAP algorithm of cells, colored by patient ID to identify residual batch effects. (N=33,038 cells from 38 patients). **(D)** As in (C), but colored by cell type assigned based on RNA expression of canonical surface markers. **(E)** As in C, but colored by 72 hours survival.

### Cell type assignment

We assigned cells in our dataset to their respective PBMC subtype based on RNA expression of canonical surface markers (supplementary **Table S2**). Previous work has indicated that the expression of genes encoding surface markers is highly correlated to protein levels of those surface markers (*7*). To verify that this was the case in this clinical context of patients under extreme physiological stress, we also analyzed these samples by conventional flow cytometry (FC). We compared the proportions of cells in the major lymphocyte subtypes as defined by the gene expression data vs. direct measurement of surface markers by FC. Cell assignments between the two methodologies were highly correlated (**Fig. 2B**).

Unsupervised clustering based on genome wide gene expression after data pre-processing (see Materials & Methods) revealed minimal clustering by patient ID (**Fig. 2C**), indicating adequate suppression of batch effects. Rather, the cells tended to cluster by major PBMC subtypes (**Fig. 2D**), based on expression of established markers of these cell populations. This observation suggests that scRNASeq can capture the major themes of cellular biology even in this clinically complex and dynamic setting—provided appropriate steps are taken to account for technical dropouts and batch effects.

### Effect of technical dropout imputation

Droplet based scRNASeq results in sparse expression data where most genes are not detected in any given cell. These “dropouts” are both biological (not all genes are actively being expressed by any given cell at any given time), and technical (the droplet based scRNASeq method, while very high throughput, will fail to detect some RNA transcripts that are in fact present in a cell). To circumvent this shortcoming, technical dropouts were imputed using the ALRA algorithm (*8*). Prior to imputation of technical dropouts, only 54% of cells could be unambiguously assigned to a specific PBMC subtype, compared to 81% after imputation. However, we wanted to verify that this imputation restored biologically meaningful information as opposed to introducing noise into the data. That is, we wanted to be sure this approach turned false negatives into true positives, as opposed to turning true negatives into false positives. To verify that this was the case, we examined the correlation with our FC data before and after imputation. Notably, imputation had no significant effect on either the slope of the regression line or the R^2^ between the proportion of lymphocytes assigned to each subclass by either FC or scRNASeq (in fact, both values increased slightly after imputation—supplemental **Table S3**). The fact that the correlation between lymphocyte subset proportions as defined by FC and scRNASeq did not deteriorate following imputation suggests that the substantial increase in cell assignment achieved by imputation identified the true biological type of these cells as opposed to random noise.

### Conventional cell types not predictive of survival

There was no clear tendency of cells to cluster based on outcome (**Fig. 2E**), indicating that transcriptional events related to survival were obscured by the dimensional reduction required to visualize gene expression in this way. We therefore proceeded to compartmentalize our analysis according to major PBMC subtypes. We sought to determine whether conventional PBMC classification could predict outcome. Using our gene expression-based cell type assignments, we looked for differences in proportions of these cell types in surviving and non-surviving patients. Although some trends were apparent, none of the observed differences were statistically significant after adjustment for multiple comparisons (**Fig. 3**). This suggested we needed to look deeper into each major cell type to try to identify novel sub-populations that may relate to survival.

**Figure 3.**
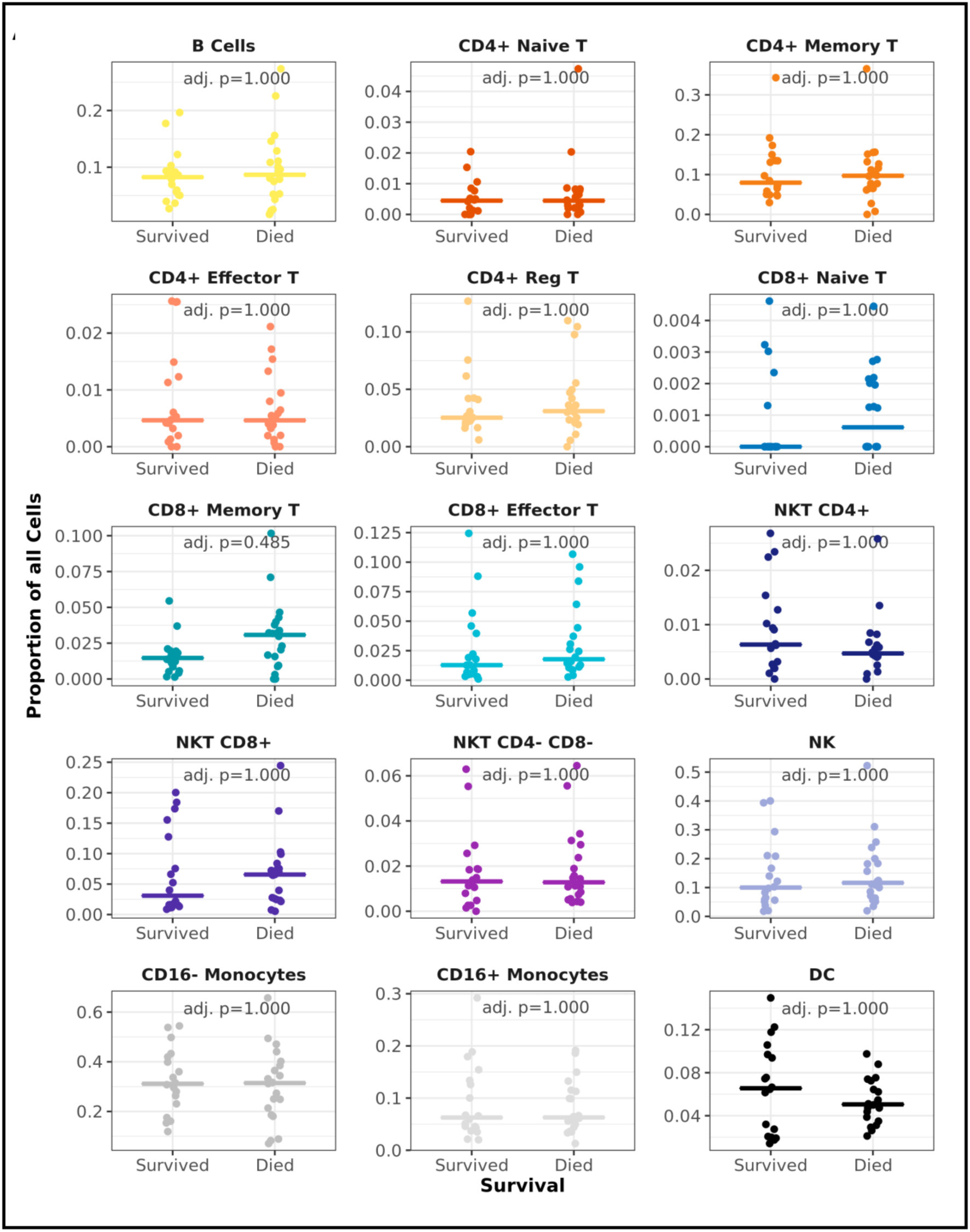
Major PBMC subtypes do not predict survival. Proportion of all PBMCs in each major PBMC subtype, stratified by 72 hour survival. Colored by cell type, matching those in Fig. 2D. P-value is for t-test, adjusted for multiple comparisons by method of Holm.

### Biological processes associated with survival

We identified the set of highly variable genes (Supplemental Data S2) within our dataset based on normalized dispersion (*9*). For each patient, we then quantified the proportion of cells of each subtype that expressed each of these genes. For each gene, the proportions among surviving patients were compared to the proportions among non-surviving patients using the Wilcoxon rank sum test. This allowed us to identify genes within each PBMC subtype that were associated with survival (blue bars in **Fig. 4A**) or non-survival (red bars in **Fig. 4A**). We clustered the genes based on this measure of differential expression, and annotated the resulting gene clusters based on their enrichment (*10*) for Gene Ontology biological process terms (**Fig. 3B**, and Supplemental **Table S4**). Within multiple PBMC subtypes, patients who died exhibited higher proportions of cells expressing genes associated with GO terms related to inflammation including antigen binding and receptor ligand binding. This suggests a widespread increase in inflammatory response across cell types in these non-surviving patients. This is consistent with the higher plasma levels of pro-inflammatory cytokines observed in non-surviving patients as noted above.

**Figure 4.**
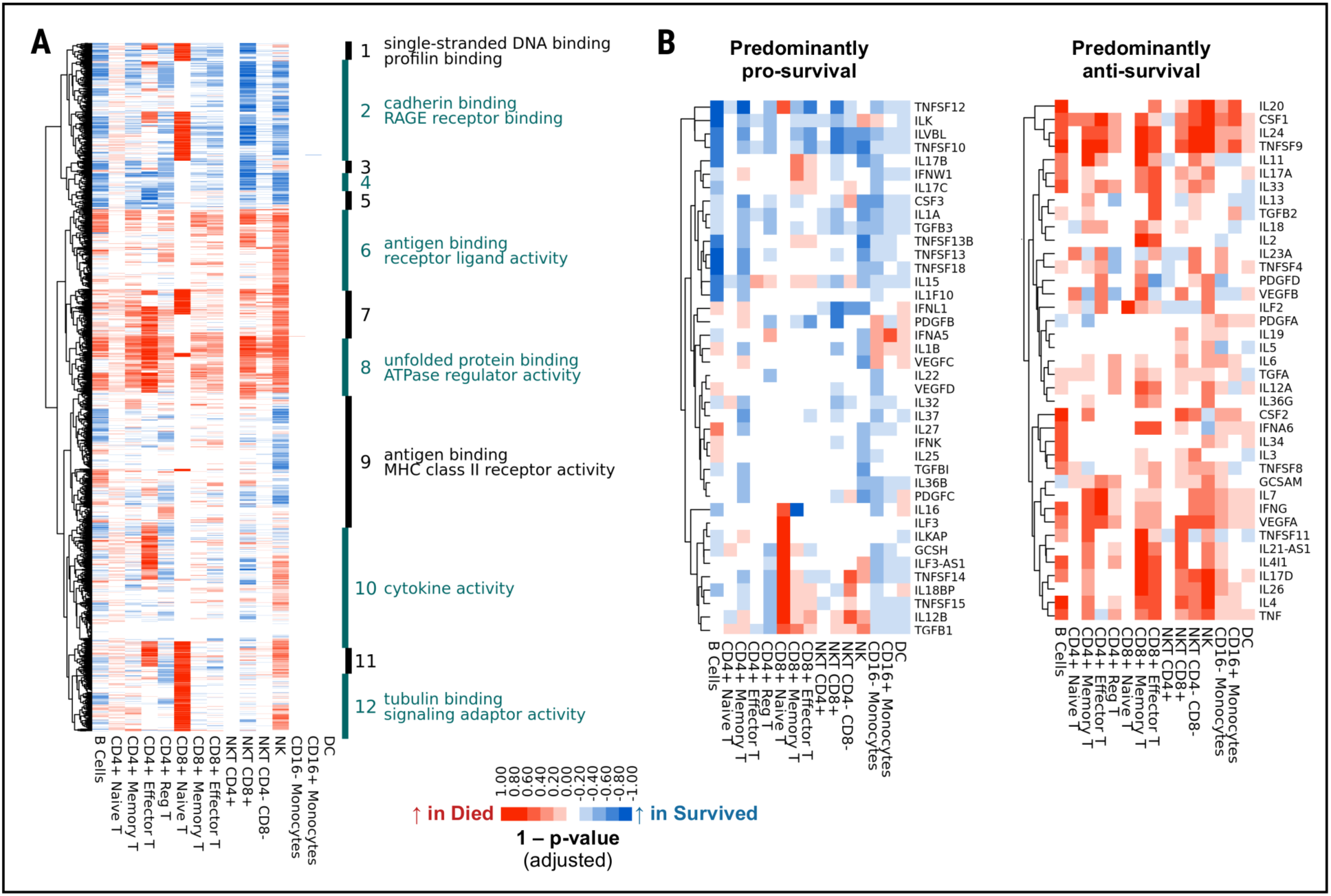
Differential gene expression and biological function analysis. Cell-type specific differential gene expression. (A) Proportions of cells expressing each highly variable gene was compared between surviving and non-surviving patients (72 hour). For each gene, the p-value (adjusted using the FDR method) for difference in proportion of cells expressing that gene between surviving and non-surviving patients (72 hour) was calculated within each PBMC subtype. Genes expressed in more cells from surviving patients than non-surviving patients within a given PBMC subtype are colored blue, while genes expressed in more cells from non-surviving patient are colored blue. The data was then clustered and plotted. The major cluster branches were analyzed for GO term enrichment (Biological Function ontology). The top two significant GO terms (where applicable) are shown. A complete list of significant GO terms is provided in the supplementary materials. (B). As in A, but restricted to cytokines.

However, other processes appeared to have more discreet patterns of activation within specific cell types. Genes associated with cadherin binding and RAGE receptor binding exhibited an inverse relationship between CD8^+^ naïve T cells compared to CD8^+^ NKT cells. Non-surviving patients exhibited higher proportions of CD8^+^ naïve T cells expressing genes related to these biological processes at the time of initiation of VA-ECLS, while the inverse was true of CD8^+^ NKT cells. Although the CD8^+^ naïve T cells constituted a small cell population, these cells can proliferate when they encounter their target antigens (*11*). Similarly, genes related to cytokine activity were more strongly upregulated in CD4^+^ effector cells and natural killer cells from non-surviving patients, whereas this trend was either not as strong or reversed in other cell types.

We also examined the expression of genes encoding cytokines themselves (**Fig. 4B**). We again saw widespread upregulation of pro-inflammatory cytokines across most cell types in non-surviving patients. This signal was particularly strong in CD8^+^ effector and memory T cells as well as natural killer and NKT cells (with the exception of CD4^+^ NKT cells).

### Novel surface markers associated with survival

The results of our analysis of plasma cytokine levels as well as gene expression patterns in specific PBMC subtypes suggested that surviving and non-surviving patients exhibit different inflammatory responses either leading to or in response to circulatory collapse and subsequent VA-ECLS. While this may be simply a manifestation of distinct disease processes, we hypothesized that there were one or more specific immune cell populations that were contributing to this difference in inflammatory response. In an effort to isolate these cells, we next focused our differential gene expression analysis on surface markers (**Fig. 5A**). Two surface markers exhibited differential expression with a false discovery rate of 10% or less. These were CD52 (overexpressed by CD8^+^ NKT cells among surviving patients), and CD36 overexpressed by CD4^+^ effector T cells among surviving patients. The CD4^+^/CD36^+^ effector cell population was very small (0.06% of all cells) compared to the CD8^+^/CD52^+^ NKT cells population (5% of all cells). The CD8^+^/CD52^+^ NKT population was selected for downstream validation.

**Figure 5:**
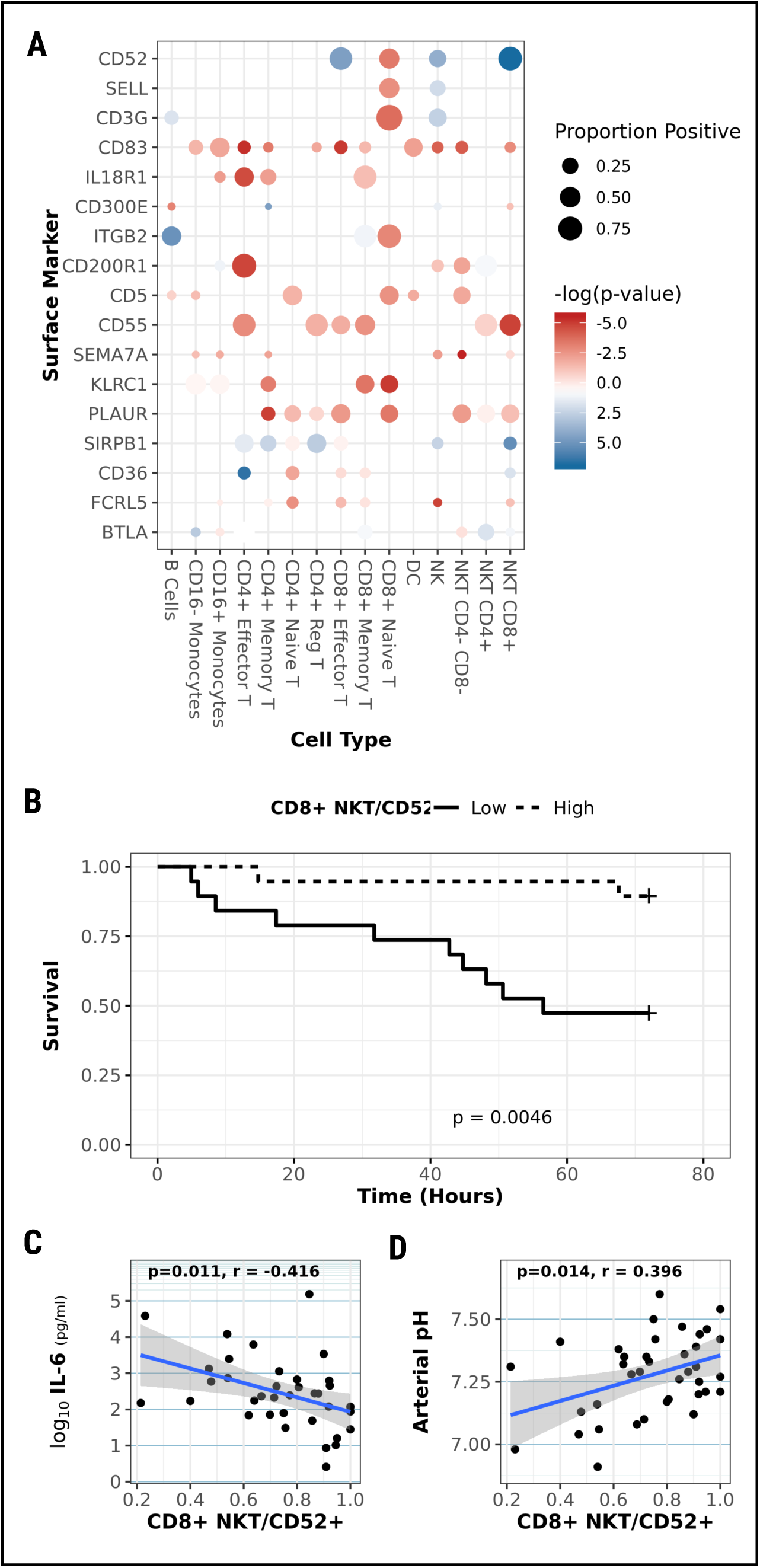
Novel cell surface marker identification. **(A)** Differential expression analysis of surface markers. Blue spots correspond to surface markers expressed by more cell from surviving patients than non-surviving patients within each cell subtype, while red spots correspond to surface markers expressed by more cells from non-surviving patients. Intensity of color corresponds to the negative log of the p-value for the test for difference in proportions (more intense color = more significant). Size of each spot is proportional to proportion of all cells of each subtype that express that marker. **(B)** 72 hour survival curves for patients with high vs. low proportion of CD8+ NKT cells that express CD52+, using the median as the threshold. P-value is for the log-rank test between groups. **(C)**. Correlation between plasma IL-6 levels and proportion of CD8+ NKT cells that are CD52+ for each patient (N=36). (D) As in C, but for arterial pH (N=38). P-values for C & D are for Pearson’s correlation coefficient.

We stratified the patients based on the proportion of their CD8^+^ NKT cells that were CD52^+^, using the median proportion as the cutoff. This stratification identified patients with significantly different mortality risks on VA-ECLS during the first 72 hours of support (**Fig. 5B)**. Patients in the low CD8^+^/CD52^+^ NKT group were more than 6 times more likely to die in the first 72 hours of VA-ECLS compared to the high CD8^+^/CD52^+^ NKT group (χ^2^ = 8.1, p=0.0043 by log rank test).

The proportion of CD8^+^ NKT cells in each patient that were positive for CD52 as determined by scRNASeq correlated inversely with plasma IL-6 levels (**Fig. 5C**). As CD8^+^/CD52^+^ NKT cells were associated with improved survival and lower levels of IL-6, we expected that this proportion would also correlate with arterial pH at start of VA-ECLS, and this proved to be the case (**Fig. 5D**).

### Validation of CD8^+^/CD52^+^ NKT Cells as predictors of outcome

The preceding analysis relied on FDR adjustment of p-values to control the family wise error rate of our identified survival markers. Based on this measure, there is less than a 10% chance that the predictive nature of CD8^+^/CD52^+^ NKT cells was due to random noise in the data. Nevertheless, it remains possible that this observation represents a biological reality for these patients that cannot be generalized to other patients due to either random variation or undetected confounding. To evaluate this possibility, we performed flow cytometric analysis of a second cohort of 21 patients that were not included in the original scRNASeq analysis. For each patient, we quantified the proportion of CD8^+^ NKT cells that were CD52^+^ (**Fig. 6A**). The same gate was used across all patients to define the CD52+ cells based on their empiric distribution (Supplemental **Fig. S1**). Using the same cutoff defined in the initial cohort, we stratified these 21 patients into “High CD52^+^” and “Low CD52^+^” based on their CD52 levels in CD8^+^ NKT cells. We compared these two groups in terms of the clinical parameters we analyzed in our initial cohort (**Fig. 6B**). Again, while there was no statistically significant difference in the age of these two groups, the High CD52^+^ patients exhibited less acidosis. Intriguingly, the High CD52^+^ patients in this validation cohort had higher SOFA scores at the start of VA-ECLS, and worse renal function. But despite these unfavorable clinical characteristics, the High CD52^+^ group once again exhibited improved survival relative to the Low CD52^+^ group (**Fig. 6C**), Patients in the low CD8^+^/CD52^+^ NKT group were nearly 5 times more likely to die in the first 72 hours of VA-ECLS compared to the high CD8^+^/CD52^+^ NKT group (χ^2^ = 4.1, p=0.043 by log rank test).

**Figure 6:**
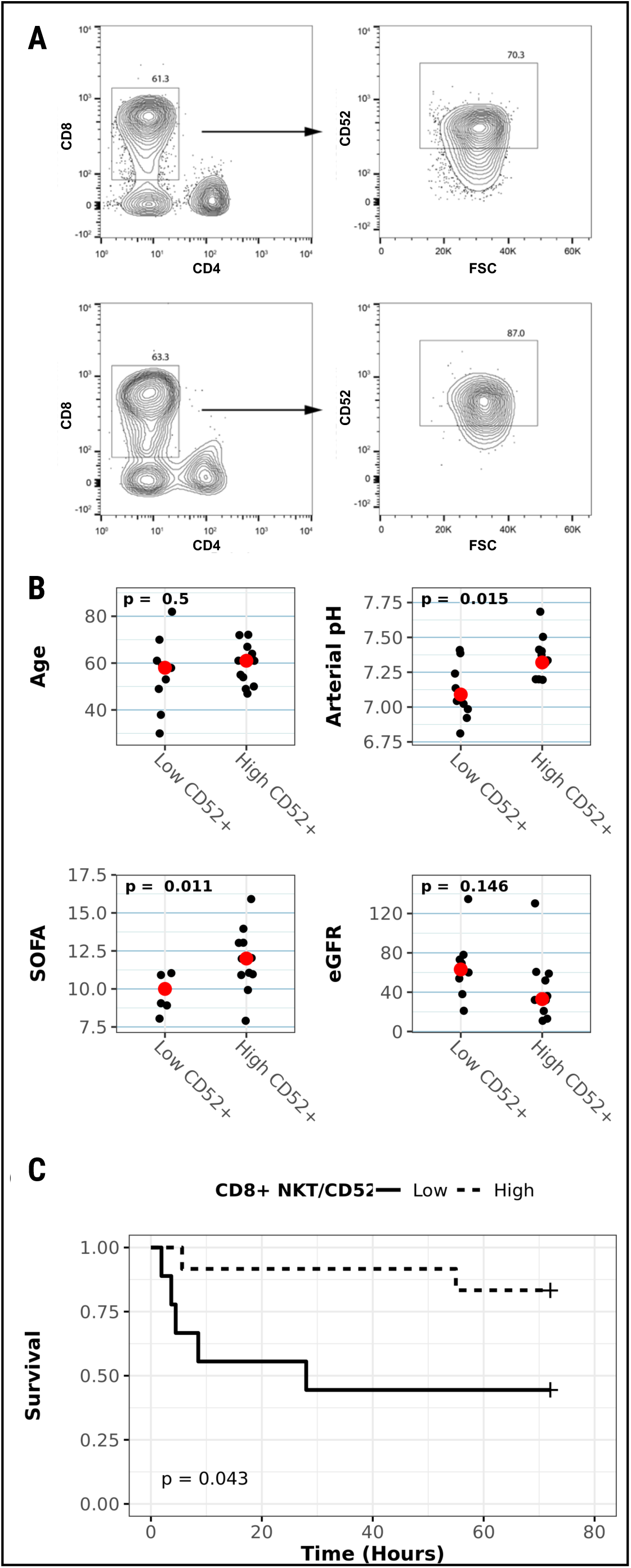
Validation of CD8+/CD52+ NKT cells as predictor in separate cohort. A second cohort of 21 VA-ECLS patients was studied by mean of conventional flow cytometry to validate our findings from the initial cohort. **(A)** Representative density plots from non-surviving (top) and surviving (bottom) patient. Left hand panels depict gating for CD8+ cells, followed by CD52 level in right hand panel. CD52 gate has been set based on distribution of CD52 across all patients (supplemental **Fig. S2**). **(B)** Key clinical parameters of patients, stratified by proportion of CD8+ NKT cells that were CD52+, using the same threshold as defined in first cohort. (N=21, p-values are for t-tests comparing the two groups.) (C). 72 hour survival curves for patients with high vs. low proportion of CD8+ NKT cells that express CD52+, using the threshold defined in the initial cohort. (N=21, p-value is for the log-rank test between groups.

## Discussion

Inflammation has long been identified as a key factor in mortality in critical care populations (*12*). With the development of new biologic agents the opportunities for a more specific modulation of the inflammatory response has grown exponentially. As a consequence, intriguing opportunities for new treatments targeting details in the inflammatory response opens up. In this study, for the first time, we combine flow cytometry, cytokine measurement, and single cell RNASeq analysis of PBMCs from patients being put on VA-ECLS due to circulatory collapse in the context of acute illness.

Single cell transcriptomics was developed over 25 years ago (*13-15*, reviewed in *16*), but the technology made a quantum advance in terms of throughput in the last several years as a result of development of microfluidics based cell encapsulation systems (*6, 9, 17*). The ability to quantify the transcriptomes of thousands of individual cells holds the promise to reveal new information about heterogeneous disease states, enabling fine tuning of personalized medicine efforts to target specific cell populations (*18*). This technology holds the promise to uncover new information about the pathophysiology of complex disease states in high detail. Initial efforts in this direction with human material have focused on cancer (*4, 19-21*), although examples in other diseases are emerging (*22, 23*) including one example of identification of a PBMC repertoire associated tuberculosis progression (*2*) and a second related to survival in acute infectious disease based on analysis of ∼100 cells from 2 patients (*3*).

While the promise of the scRNASeq approach is substantial with respect to deeper understanding of pathological processes, the extent to which scRNASeq is both feasible and informative in the dynamic setting of critical illness remains largely unknown. Our results provide evidence that not only can this technology detect biological signal in a heterogeneous and rapidly changing clinical context but can do so in a way that reveals deeper understanding of physiologic events associated with—and potentially driving—clinical outcomes.

VA-ECLS is a powerful stimulator of the immune response against a background of already tenuous perfusion and end organ function (*24*). Whether this immune response is adaptive or mal-adaptive remains unclear. On the one hand, immunoparalysis was associated with worse outcomes (though not in a statistically significant fashion) in one small series (*25*). Also, the 2006 ARDS Network Trial (*26*) found worse outcomes when steroids were started late in the disease course, and no change in outcomes when they were started earlier (despite short term improvements in physiologic measures of ventilation and perfusion). On the other hand, the use of steroids has been associated with survival in elderly patients in one registry based review (*27*) as well as dramatic clinical improvement on VA-ECLS in a number of case reports (*28-31*). These contradictory results underscore the need for delineating the inflammatory response in these patients in more detail. Furthermore, all of this evidence comes from the setting of acute respiratory failure. There is little or no data to inform us about the role of inflammation with respect to survival among patients undergoing VA-ECLS for acute decompensated heart failure. In this study we show that the complexity of the inflammatory response in patients with circulatory collapse can be deconvoluted by means of scRNASeq combined with appropriate bioinformatic methods and validation with flow cytometry.

We found that higher proportions of CD8^+^ NKT cells that were CD52^+^ was associated with improved survival. CD52 has been implicated as an immunomodulatory protein along several lines. CD4^+^CD25^+^FoxP3 cells suppress T-cell functions through the binding of free CD52 of Siglec-10 on CD52^-^ T-cells. Subsequently it was shown that cross linking of CD4^+^/CD52^+^ T-cells by the 4C8 antibody leads to their expansion and resultant suppression of other CD4^+^ and CD8^+^ cells. Notably, these cells suppress IL-2 production by other T populations upon subsequent stimulation, and prevented lethal graft-versus-host reactions in SCID mice (*32, 33*). While this data all focuses on CD4+/CD52+ T cells, additional data suggests there is a subset of CD8^+^ cells that are protective against autoimmune disease (*34*), and that these protective cells are likewise CD52^+^ (*35*).

In our study the strongest association with survival among CD52^+^ cells was among CD8+ NKT cells and to a lesser degrees CD8^+^ effector T-cells. The association was confirmed both on the mRNA and protein level across two separate cohorts. Since the patients were sampled for cells around the time of cannulation, it is unlikely that the CD52+ population is a response to VA-ECLS itself. Rather, this population may reflect the degree to which the immune system is poised for a pathological vs. adaptive response to the inflammatory stimulation of VA-ECLS. Insofar as these cells promote a permissive (rather than reactive) immune state, this observation is consistent with the hypothesis that attenuating the immune response to VA-ECLS may be beneficial in patients with severe heart failure. Moreover, the strong negative association of these cells to the tissue-aggressive inflammatory marker IL-6—which was itself associated with decreased survival—supports the immuno-attenuating effect of these cells.

Fluidics based scRNASeq enables transcriptomic characterization of individual patient cells at high throughput. The resulting data presents specific challenges and pitfalls. However, we have demonstrated that these factors can be overcome through appropriate analysis techniques and that this technology allows detection of biologically meaningful signal in these cells even in a dynamic and complex clinical setting. We anticipate that future applications of this approach will continue to reveal new information about the roles of both known and novel cell populations in human disease, and this information will continue to establish new biomarkers and therapies to the benefit of our patients.

Even if our current results do not allow us to determine whether the associations identified are cause or effect, they do delineate important pathways and interesting drug targets for future interventional studies. Subsequent studies examining the effect of activation of CD8+/CD52+ NKT cells via cross linking with 4C8 in animal models of circulatory collapse will be an important next step to clinical application of this strategy.

## Materials and Methods

### Patient materials

This study was approved by our institution’s Institutional Review Board, and all patients or their representatives gave informed consent to participate. All samples and data were anonymized prior to the analysis described here. Plasma albumin, AST, ALT, bilirubin, pH, lactate, creatinine, sodium, potassium, magnesium, and glucose were measured in our clinical laboratory as part of standard of care evaluation of the patients.

Approximately 10ml of whole blood from VA-ECLS patients at time of boarding (mean difference from pump start time: ±79 minutes) were collected in EDTA Vacutainers. Peripheral blood mononuclear cells (PBMC) were isolated by density centrifugation using Ficoll-Paque PLUS (GE Healthcare) and cryopreserved in CryoStor (Sigma-Aldrich) at approximately 4×10^6^ cells per vial.

### Cytokine Measurement

Approximately 10ml of whole blood from ECLS patients at time of boarding (mean difference from pump start time: + 79 minutes) were collected in EDTA Vacutainers. Plasma was isolated by centrifugation. The concentrations of cytokines in plasma were measured using the Bio-Plex Pro™ Human Cytokine 17-plex Assay kit immunoassay run on the Bio-Plex 200 (BioRad), with reference to an 8 point standard curve.

### Flow cytometry

Cryopreserved PBMCs were rapidly thawed at 37°C, washed in RPMI (10% FBS, 2mM L-glutamine, 1:10000 Benzonase), and rested overnight in 200µl of RPMI (10% FBS, 2mM L-glutamine) in a 96 well plate at a concentration of 1.0×10^6^ cells per well. Prior to surface staining non-viable cells were labelled with Fixable Viability Stain 450 (FVS450) and Fc receptors were blocked. A multiparameter flow cytometry panel was designed for detection of surface antigens. The panel consisted of directly conjugated anti-human antibodies; CD3-BB515, CD4-BUV395, CD8-APC-H7, CD19-APC, CD56-APC-R700, and CD16-BV510. CD52 expression was measured by CD52-PE-4C8 directly conjugated anti-human antibodies (BD Life Sciences). For cell surface markers, cells were stained in PBS supplemented with 2% FBS and 2mM EDTA for 35 minutes at 4°C. Stained cells were analyzed on a BD Influx flow cytometer equipped with 488nm, 355nm, 561nm, 405nm, and 640nm lasers, using a 100µM nozzle, at 20 psi, and an offset of 1.0. Gating strategy for major PBMC populations is summarized in **Fig. S2**. Gating strategy for the CD52 analysis in the validation cohort is summarized in supplemental **Fig. S1**. Flow experiments included single-stained controls, fluorescence minus one controls, and well-characterized healthy control. Acquisition of flow cytometry data was performed using Sortware v1.2 and analyzed with FlowJo v10.0.8.

### Single cell encapsulation and reverse transcription

At the time of cell encapsulation for single cell RNASeq, cryopreserved PBMCs were rapidly thawed at 37°C, washed in twice in RPMI (10% FBS, 2mM L-glutamine, 1:10000 Benzonase) and 2×10^5^ cells were resuspended in 1x PBS. To exclude non-viable cells from sorting, 3 nM of SYTOX Green (Thermo Fisher Scientific) was added to each sample tube. Cells were sorted on a BD Influx flow cytometer using a 100µM nozzle, at 20 psi, and an offset of 1.0. The following gating hierarchy was used: PBMCs were separated from debris based on distribution of light scatter by SSC/FSC; cell doublets were excluded by signal pulse characteristics of FSC-W/FSC-H and SSC-H/SSC-A. Viable cells with intact cell membranes were gated as SYTOX Green negative. For each patient sample 20,000 events were sorted (51.2ul) directly in 112.8µl of 1xPBS. Prior to InDrop 36ul of optiprep was added to each tube.

Thawed, sorted and diluted cells were encapsulated along with barcoded beads and reverse transcription reagents using the inDrop platform (1CellBio, Watertown, MA). Flow rates were adjusted periodically throughout the experiment, with the help of high speed video microscopy, to ensure that the number of droplets containing one bead was maximized while minimizing droplets with two or more beads. Run times were calculated to capture 1500 cells per sample. Each sample was run on a separate, freshly silanized microfluidics device. Reverse transcription was performed following the manufacturer’s protocol. Briefly, barcoding oligo were cleaved by exposing each droplet emulsion aliquot to UV light for 10 minutes. The emulsions were then incubated at 50°C for one hour, and then 70°C for 15 minutes. The emulsion was then broken, and the aqueous phase containing the cDNA removed. The cDNA was cleaned up with MinElute columns (Qiagen, Hilden, Germany) and excess barcodes enzymatically removed. Second strand synthesis was performed using the NEBNext Ultra II second strand synthesis kit (NEB, Ipswich, MA) according to the manufacturer’s protocol. The samples were then again cleaned up on MinElute columns, and sample integrity confirmed by Bioanalyzer (Agilent, Santa Clara, CA). *In vitro* transcription was then performed using the NEB HiScribe High Yield RNA synthesis kit according to the manufacturer’s instructions. Sample integrity was again verified by Bioanalyzer. The IVT products were then reverse transcribed using random hexamers. Amplification cycles were optimized by diagnostic qPCR, and then the samples were amplified using unique PE1/PE2 indexing primers such that samples could be pooled prior to sequencing. Amplified cDNA was then cleaned up using AMPure beads (Beckman Coulter, Indianapolis, IN). Library integrity and fragment size was confirmed by BioAnalyzer prior to sequencing.

### Data and code availability

The single cell RNASeq expression data (as a matrix of raw counts) and supporting metadata is available from GEO (GSE127221). Code used to perform the analysis and generate the figures, with accompanying documentation and explanation including system requirements and dependencies, is available from Github at http://github.com/vanandelinstitute/va_ecls. A rendered html file providing the analysis details is also provided along with this manuscript as Supplmental Data File S3.

### Sequencing

Prepared libraries were normalized and pooled, and sequenced on a NovaSeq 6000 sequencer (Illumina, San Diego, CA) using the S2 100 cycle kit. Read one was 36 cycles, the index read was 6 cycles, and Read 2 was 50 cycles. Cells were sequenced to an approximate depth of 90,000 reads per cell. Resulting sequencing data was converted to demultiplexed FASTQ files prior to downstream analysis.

### Data processing

The sequencing data was aligned to the human genome (assembly GRCh38) and unique feature counts obtained using the software pipeline developed by the inDrop manufacturer (https://github.com/indrops/indrops). The raw count data was then filtered, normalized, imputed, and batch corrected using tools for the R statistical analysis platform. Full details of the data processing and analysis are presented in Supplemental datafile D1, and is also available as an R markdown document at https://github.com/vanandelinstitute/va_ecls.

### Statistical analysis

For differential gene expression analysis, the expression matrix was filtered to include only variably expressed genes as described (*36*). Briefly, for each gene, the mean was calculated across all cells. The dispersion of (variance / mean) was also calculated for each gene across all cells. The genes were then split into 20 bins based on mean expression. Within each bin, dispersions were converted to robust z-scores (the absolute difference between each dispersion and the median dispersion for that bin, divided by the median absolute deviation for that bin). Genes with a dispersion z-score > 2.0 were retained for further analysis.

Given that the single cell RNASeq expression data was a sparse matrix, we compared patients in terms of proportions of cells expressing genes of interest. For any given gene, the proportion of cells (of a given subtype of interest) was calculated. Surviving patient and non-surviving patients were then compared in terms of median proportion, and difference between patient groups was tested by mean of Wilcoxon rank sum analysis of the proportions in each group. Given the large number of genes under analysis, all p-values were adjusted using the false discovery rate method (*37*).

When comparing proportions of cells in each of the major subtypes between outcome groups (**Fig. 4A**), the same approach was used, but since the number of comparisons was relatively small and we wanted to avoid any type I errors (as opposed to simply constraining the family wise error rate), p-values were adjusted using the method of Holm.

Survival analysis was performed by plotting Kaplan Meijer curves and comparing the curves using the log-rank test. Given that only 3 survival curves were analyzed, no p-value adjustment was performed (although the results would have remained significant even under the most stringent adjustment including Bonferroni correction).

## Supporting information

Analysis details and code

Variably expressed genes

## Supplementary Materials

Tables S1-S5 and Figures S1-S2, and supplemental data files D1 and D2 are provided in separated files.

## Acknowledgments

We would like to acknowledge Jennifer Schuitema for overseeing recruitment, consent, and collection of blood samples, the ECMO nursing staff for collecting blood samples and processing plasma after-hours, and David Chesla and Donald Daley from the Spectrum Health Biorepository for sample processing during after-hours. The authors thank the Van Andel Genomics Core for providing sequencing facilities and services.

## Author contributions

EJK and SJ conceptualized and conceived the project. EJK also performed the analysis, developed methodology, and wrote the manuscript. MW performed the flow cytometry work described in this work, analyzed data, and reviewed and edited the manuscript. HM, and EE performed the single cell encapsulation, and reviewed and edited the manuscript. CK assisted with patient enrollment and data collection. EG, ML, SF, TB, GM, TT, MD, and PW provided clinical resources and supervision for the project and reviewed the manuscript. NMS performed analysis of clinical data. SJ also acquired funding, developed methodology, supervised the project, and wrote the manuscript.

## Data and materials availability

The expression data is available as a matrix of raw counts from GEO (GSE127221). Code used to perform the analysis and generate the figures is available from Github at http://github.com/vanandelinstitute/va_VA-ECLS.

