## Supplementary material for "The inflammasome of circulatory collapse: single cell analysis of survival on extracorporeal life support": Analysis details and code

### Data Analysis Supplement

*March 6, 2020*

#### Introduction

This document was prepared in support of our paper:

The inflammasome of circulatory collapse: single cell analysis of survival on extra corporeal life support.

Eric J. Kort MD, Matthew Weiland, Edgars Grins MD, Emily Eugster MS, Hsiao-yun Milliron PhD, Catherine Kelty MS, Nabin Manandhar Shrestha PhD, Tomasz Timek MD, Marzia Leacche MD, Stephen J Fitch MD, Theodore J Boeve MD, Greg Marco MD, Michael Dickinson MD, Penny Wilton MD, Stefan Jovinge MD PhD

The following sections document how the data for this paper were processed and how the figures were generated. Those who wish to do so can recreate the figures from the paper from data posted on GEO under accession GSE127221, and the code below. For efficient compiling of this document, the "eval=FALSE" option was set globally. Readers desiring to repeat the analysis can either run each code chunk below manually, or change the setting line at the top of the Rmd file from:

```
knitr::opts_chunk$set(echo = TRUE, eval=FALSE)
```

to

```
knitr::opts_chunk$set(echo = TRUE, eval=TRUE)
```

And then knit the entire document, a process which may take several hours, and may also fail if insufficient RAM is available.

The result of running this code is that all processed data and generated figures will be produced and saved to the Final_Data directory. The figures should match exactly what is in the publication (except for small variations due to random processes–for example, the jitter applied to do plots to enable visualization of overlapping points may be slightly different).

Since the processed data elements created below are save as RDS files in the chunks that create them, you can execute just some of the chunks and then pick up where you left off later.

#### Pre-requisites

Running the following analysis requires a computer with R version >= 3.5.0 and 128GB of RAM, with `pandoc`, and development libraries for SSL, XML, and curl installed. This would be achieved on a debian flavored server as follows:

```
sudo -s
apt-get update
apt-get -y install pandoc
apt-get -y install libssl-dev
apt-get -y install libcurl4-openssl-dev
apt-get -y install libxml2-dev
```

To run the following analysis, the file `GSE127221_PBMC_merged_filtered_recoded.rds` must be obtained from the GEO series related to this study (GSE127221). The second panel of figure 4b requires the list of genes from Supplmental Table S5, which is available in the gitub repository as `Final_Data/table_s5.txt`. The following code assumes these files are within a subdirectory named "Final_Data" underneath the working directory. Note that cloning the github repo will create the necessary directory structure with supporting files. However, `PBMC_merged_filtered_recoded.rds` is too large for the gitub repository, and thus must be obtained from GEO using the linke above and saved in the Final_Data directory.

For example:

```
git clone https://github.com/vanandelinstitute/va_ecls
cd va_ecls/Final_Data
wget ftp://ftp.ncbi.nlm.nih.gov/geo/series/GSE127nnn/GSE127221/suppl/GSE127221_PBMC_merged_filtered_rec
gunzip *.gz
cd ..
```

If you are not using Rstudio, the following step (from within an R session) is required to install the rmarkdown tools. If you are using Rstudio, you can skip this.

```
install.packages("rmarkdown")
# the next line will render this documnt, generating all the
# intermediate data files and figures
rmarkdown::render("final_analysis.rmd")
```

If you are using Rstudio, just click the Knit button, or run each chunk individually.

Next we check for, conditionally install, and load the required libraries:

```
setRepositories(ind = c(1, 2, 3, 4))
reqpkg <- c("clusterProfiler", "cowplot", "dendextend", "devtools",
    "doParallel", "DOSE", "dplyr", "foreach", "ggplot2", "GGally",
    "ggpubr", "ggplotify", "grid", "irlba", "Matrix", "org.Hs.eg.db",
    "pheatmap", "reshape2", "rsvd", "survminer", "stringr", "survival")

missingpkg <- reqpkg[-which(reqpkg %in% installed.packages())]
if (length(missingpkg)) install.packages(missingpkg)

if (!"harmony" %in% installed.packages()) devtools::install_github("immunogenomics/harmony")
if (!"uwot" %in% installed.packages()) devtools::install_github("jlmelville/uwot")
if (!"rstatix" %in% installed.packages()) devtools::install_github("kassambara/rstatix")
```

```
reqpkg <- c(reqpkg, "harmony", "uwot", "rstatix")
for (i in reqpkg) library(i, character.only = TRUE)
```

#### Data Pre Processing

The data file `GSE127221_PBMC_merged_filtered_recoded.rds` contaings sequencing data that was aligned and converted to UMI counts with the inDrop pipeline. Barcodes were filtered to retain only cells with at least 500 unique counts.

As shown below, we then further filtered this dataset to select cells with between 750 and 7500 unique counts in an effort to select true, single cells. We then normalized by library size, scaled by a constant (10000), and log transformed, as follows:

```
dat <- readRDS("Final_Data/GSE127221_PBMC_merged_filtered_recoded.rds")

# filtering, somewhat subjectively, for true, single cells
cellCounts <- Matrix::rowSums(dat)
dat <- dat[which(cellCounts < 7500 & cellCounts > 750), ]
geneCounts <- Matrix::colSums(dat)
dat <- dat[, which(geneCounts >= 10)]

# we want cells as row and genes as columns for ALRA
dat.m <- as(dat, "matrix")
rm(dat)

# normalize to library size and log transform
sizes <- rowSums(dat.m)
dat.n <- sweep(dat.m, MARGIN = 1, sizes, "/")
dat.n <- dat.n * 10000
dat.n <- log(dat.n + 1)
saveRDS(dat.n, file = "Final_Data/PBMC_merged_filtered_normalized.rds")

source("https://raw.githubusercontent.com/KlugerLab/ALRA/master/alra.R")

# ALRA uses the random SVD for performance reasons.  If we
# don't set a seed, the results of the imputation will be
# very nearly but not exactly the same between runs.
set.seed(1010)
dat.imp <- alra(dat.n, k = 50, q = 40)[[3]]
rownames(dat.imp) <- rownames(dat.n)
saveRDS(dat.imp, file = "Final_Data/PBMC_merged_filtered_alra.rds")
```

For batch effect removal and UMAP visualization, we first need the principle component loadings for the imputed dataset. Calculating just a partial PC matrix (the first 20 PCs, which is plenty as we shall see from plotting the standard deviations for each PC) using the `irlba` package makes this task much more tractable in terms of both memory and time requirements.

```
library(irlba)
dat.imp <- readRDS("Final_Data/PBMC_merged_filtered_alra.rds")
dat.pc <- prcomp_irlba(dat.imp, n = 20)
rownames(dat.pc$x) <- rownames(dat.imp)
plot(dat.pc$sdev)
saveRDS(dat.pc, file = "Final_Data/PBMC_merged_filtered_alra_pc.rds")
```

Next we regress away donor (batch) effect using the Harmony algorithm.

```
library(harmony)

dat.pc <- readRDS("Final_Data/PBMC_merged_filtered_alra_pc.rds")

# patient id is the library id and each patient was a
# separate sequencing library
id <- gsub("\\..*", "", rownames(dat.pc$x))

dat.pc.h <- HarmonyMatrix(dat.pc$x, do_pca = FALSE, data.frame(id = id),
    "id", theta = 4)
saveRDS(dat.pc.h, file = "Final_Data/PBMC_merged_filtered_alra_pc_harmony.rds")
```

We can use these PC loadings to further reduce dimensionality (to 2 dimensions suitable for visualization) with UMAP.

```
library(uwot)

dat.pc.h <- readRDS(file = "Final_Data/PBMC_merged_filtered_alra_pc_harmony.rds")
set.seed(1010)
umap.h <- umap(dat.pc.h, min_dist = 0.2, n_neighbors = 15, n_threads = 7)
umap.h <- data.frame(id = gsub("\\..*", "", rownames(dat.pc.h)),
    UMAP1 = umap.h[, 1], UMAP2 = umap.h[, 2])
saveRDS(umap.h, "Final_Data/PBMC_merged_filtered_alra_pc_harmony_umap.rds")
```

Note that we could put all of this data into a `SingleCellExperiment` object or a Seurat or Monocol object. For most of the operations below which rely on the normalized count data itself, there seemed to be negligible benefit for doing so for the purposes of this analysis, so we prefered to keep the expression matrix and metadata as separate objects and work on them directly. But feel free to select another data structure that best fits your needs.

#### Plot formats

We define a theme to tweak plotting formatting (I prefer bold axis labels, etc.) and keep things consistent. We also create a helper function for the UMAP plots.

```
my_theme <- theme_bw() + theme(strip.background = element_blank(),
    strip.text = element_text(face = "bold", margin = margin(0,
        0, 5, 0), size = 10), panel.spacing = unit(1, "lines"),
    plot.margin = unit(c(0.5, 0.5, 0.5, 0.5), "cm"), plot.title = element_text(size = 10,
        margin = margin(0, 0, 5, 0), face = "bold"), axis.text = element_text(size = 10),
    axis.title.y = element_text(face = "bold", margin = margin(t = 0,
        r = 0, b = 0, l = 0, unit = "pt")), axis.title.x = element_text(face = "bold",
        margin = margin(t = 2, r = 0, b = 0, l = 0)))

my_theme_wide_lab <- theme(axis.title.y = element_text(face = "bold",
    margin = margin(t = 0, r = 10, b = 0, l = 0)))

my_theme_no_labs <- my_theme + theme(axis.title = element_text(size = 0))

my_theme_no_space <- my_theme + theme(plot.margin = unit(c(0,
```

```r
    0, 0, 0), "cm"))

my_theme_wider_lab <- theme(axis.title.y = element_text(face = "bold",
    margin = margin(t = 0, r = 20, b = 0, l = 0)))

my_theme_pretty_grid <- theme(panel.grid.major.y = element_line(color = "#8DBCD2",
    size = 0.3), panel.grid.minor.y = element_line(color = "#A8CFD1",
    size = 0.1))

# pretty print p values
fp <- function(p) {
    if (p < 0.001)
        return("p < 0.001")
    return(paste("p = ", round(p, 3)))
}

plotUmap <- function(p1, p2, size = 1, alpha = 0.1, color, label,
    palette, legend.position = "none") {
    dat <- melt(data.frame(UMAP1 = p1, UMAP2 = p2, color = color),
        id.vars = c("UMAP1", "UMAP2", "color"))
    ggplot(dat, aes(x = UMAP2, y = UMAP1, color = color)) + geom_point(alpha = alpha,
        pch = 19, size = size, stroke = 0) + annotate("text",
        x = -10, y = -12, size = 3, hjust = 0, label = label) +
        theme_bw() + my_theme + ylim(c(-12, 12)) + xlim(c(-12,
        12)) + scale_color_manual(values = palette) + theme(legend.position = legend.position) +
        guides(color = guide_legend(override.aes = list(size = 5,
            alpha = 1)))

}

# helper function to load up data (unless, in the case of the
# largeish dat.imp object, it is already loaded). This is
# only every necessary if we are running just portions of the
# code below as opposed to rendering everything in one shot
# top to bottom.

load_init <- function() {
    if (!exists("dat.imp"))
        dat.imp <<- readRDS("Final_Data/PBMC_merged_filtered_alra.rds")
    md <<- readRDS("Final_Data/metadata_clin_cyt.rds")
    ix <- which(md$group == "Initial")
    md <<- md[ix, ]
    md$Survived <<- factor(md$surv_time < 72, labels = c("Survived",
        "Died"))
}

types <- readRDS("Final_Data/cell_types.rds")
```

#### Figure 1

```r
# load our data if we haven't already
load_init()
```

```r
pv <- foreach(i = c(1:4)) %do% {
    t.test(md[, i] ~ md$Survived)$p.value
}

pv.ev <- foreach(i = c(1:4)) %do% {
    t.test(md[, i] ~ md$Survived, var.equal = TRUE)$p.value
}

# SOFA and eGFR have equal variance, the other two variables
# do not.

pv[[3]] <- pv.ev[[3]]
pv[[4]] <- pv.ev[[4]]

# extract partial dataset to clean up labels, etc.
dat <- md[, 1:4]
dat$Survived <- md$Survived
colnames(dat) <- c("Age", "Arterial pH", "SOFA", "eGFR", "Survived")

# Panel 1A

pl <- foreach(i = 1:4) %do% {
    gd <- data.frame(Survived = dat$Survived, y = dat[, i])
    ggplot(gd, aes(x = Survived, y = y)) + geom_jitter(width = 0.1,
        height = 0.1) + stat_summary(fun.y = median, color = "red",
        geom = "point", aes(group = 1), size = 3, show.legend = FALSE) +
        ylab(colnames(dat)[i]) + annotate("text", size = 3, fontface = 2,
        label = fp(as.numeric(pv[i])), x = -Inf, y = Inf, hjust = -0.1,
        vjust = 1.5) + scale_y_continuous(expand = expand_scale(mult = c(0.1,
        0.2), add = 0)) + xlab("") + theme_bw() + my_theme +
        my_theme_wide_lab + my_theme_pretty_grid + theme(axis.text.x = element_text(angle = 315,
        hjust = 0.1, vjust = 0.5)) + theme(plot.margin = unit(c(0.1,
        1, 0.2, 0.2), "lines"))
}

png("fig_1a.png", width = 1500, height = 1500, res = 300)
plot_grid(plotlist = pl, ncol = 2, align = "v")
dev.off()

# Panel 1B

pv <- foreach(i = 9:13) %do% {
    wilcox.test(md[, i] ~ md$Survived)$p.value
}

# note: we cannot do the p-value adjustment here because the
# other cytokines are not included in the dataset provided in
# this repository. So we will simply hard code the Holm
# corrected p-values here.

# pv <- p.adjust(pv, 'holm')
pv <- c(0.016174682, 0.011424845, 0.001272499, 0.008249142, 0.011424845)
```

```
# extra dataset to clean up labels, etc.
dat <- md[, 9:13]
dat$Survived <- md$Survived
colnames(dat) <- gsub("(.{2,3})_(.{1,2})_.*", "\\1-\\2", colnames(dat))

options(scipen = 1e+06)
pl <- foreach(i = 1:5) %do% {
    gd <- data.frame(Survived = dat$Survived, y = dat[, i] +
        1)
    ggplot(gd, aes(x = Survived, y = y)) + geom_jitter(width = 0.05,
        height = 0.05) + stat_summary(fun.y = median, color = "red",
        geom = "point", aes(group = 1), size = 3, show.legend = FALSE) +
        ylab(bquote(log[10] ~ bold(.(colnames(dat)[i])) ~ scriptstyle("(pg/ml)"))) +
        annotate("text", size = 3, fontface = 2, label = paste("adj.",
            fp(as.numeric(pv[i]))), x = -Inf, y = Inf, hjust = -0.1,
            vjust = 1.5) + scale_y_continuous(minor_breaks = scales::extended_breaks(15),
        trans = "log10", breaks = c(0, 1, 10, 100, 1000, 10000,
            1e+05), labels = function(x) {
            format(round(log10(x)), scientific = FALSE)
        }, expand = expand_scale(mult = c(0.1, 0.2), add = 0)) +
        xlab("") + theme_bw() + my_theme + my_theme_wide_lab +
        my_theme_pretty_grid + theme(axis.text.x = element_text(angle = 315,
        hjust = 0.1, vjust = 0.5)) + theme(plot.margin = unit(c(0.1,
        1, 0, 0.2), "lines"))
}

png("fig_1b.png", width = 1500, height = 2200, res = 300)
plot_grid(plotlist = c(pl), ncol = 2, align = "v")
dev.off()
```

#### Figure 2

Figure 2A is just a schematic providing an overview of the study. The remaining panels require inferred cell types, so we will do that first. Cell types are defined based on RNA expression of canonical surface markers (see supplemental Table S2 accompanying the paper). The original CITE-Seq paper (Zheng, et al.) showed excellent correspondence between mRNA levels and cell surface protein levels, so we decided that simply using mRNA expression as a proxy for cell surface marker protein level might be a reasonable approach. We also tried a more sophisticated anchor transfer method to leverage more the gene expression data available here, but that method performed poorly in terms of correlation to our FACS data. Since this simple method performed well, we just stuck with that.

First we define a list structure containig our cell type definitions. The "gate" field indicates whether the corresponding marker should be expressed (`gate = 1`) or not expressed (`gate = 0`). If `gate` is $> 1$, then (`gate - 1`) is used as the threshold, although this was case only applies to erythrocytes since hemolysis results in some HBA1 RNA is most droplets. We then apply these gates to each cell. Where this results in more than one cell type assignment (i.e., ambiguous assignment), the cells is assigned the "Unknown" cell type.

```
# load our data if we haven't already
load_init()

cellTypeDefs <- list(
  "B Cells" = list(markers = c("CD19", "CD3E"),
```

```r
                     gate = c(1, 0)),

  # CD4 subpopulations can by CD2+/- and FOXP3 +/-, but not double positive
  # Except for CD4 regulatory T which are double positive
  "CD4+ Naive T" = list(markers = c("CD3E", "CD4", "CD8A", "CD2", "FOXP3", "NCAM1"),
                     gate = c(1, 1, 0, 0, 0, 0)),
  "CD4+ Naive T" = list(markers = c("CD3E", "CD4", "CD8A", "CD2", "IL2RA", "FOXP3", "NCAM1" ),
                     gate = c(1, 1, 0, 0, 0, 1, 0)),
  "CD4+ Memory T" = list(markers = c("CD3E", "CD4", "CD8A", "CD2", "B3GAT1", "FOXP3", "NCAM1"),
                     gate = c(1, 1, 0, 1, 0, 0, 0)),
  "CD4+ Memory T" = list(markers = c("CD3E"
                                   , "CD4", "CD8A", "CD2", "B3GAT1", "IL2RA", "FOXP3", "NCAM1"),
                     gate = c(1, 1, 0, 1, 0, 0, 1, 0)),
  "CD4+ Effector T" = list(markers = c("CD3E", "CD4", "CD8A", "CD2", "B3GAT1", "FOXP3", "NCAM1"),
                     gate = c(1, 1, 0, 1, 1, 0, 0)),
  "CD4+ Effector T" = list(markers = c("CD3E", "CD4", "CD8A", "CD2", "B3GAT1", "IL2RA", "FOXP3", "NCAM1
                     gate = c(1, 1, 0, 1, 1, 0, 1, 0)),
  "CD4+ Reg T" = list(markers = c("CD3E", "CD4", "CD8A", "IL2RA", "FOXP3", "NCAM1"),
                     gate = c(1, 1, 0, 1, 1, 0)),

  "CD8+ Memory T" = list(markers = c("CD3E", "CD4", "CD8A", "CD2", "B3GAT1", "NCAM1"),
                     gate = c(1, 0, 1, 1, 0, 0)),
  "CD8+ Naive T" = list(markers = c("CD3E", "CD4", "CD8A", "CD2", "NCAM1"),
                     gate = c(1, 0, 1, 0, 0)),
  "CD8+ Effector T" = list(markers = c("CD3E", "CD4", "CD8A", "CD2", "B3GAT1", "NCAM1"),
                     gate = c(1, 0, 1, 1, 1, 0)),

  "NKT CD4+" = list(markers = c("CD3E", "CD4", "CD8A", "CD19", "NCAM1"),
                     gate = c(1, 1, 0, 0, 1)),

  "NKT CD8+" = list(markers = c("CD3E", "CD4", "CD8A", "CD19", "NCAM1"),
                     gate = c(1, 0, 1, 0, 1)),

  "NKT CD4- CD8-" = list(markers = c("CD3E", "CD4", "CD8A", "CD19", "NCAM1"),
                     gate = c(1, 0, 0, 0, 1)),

  "NK" = list(markers = c("CD3E", "CD19", "NCAM1"),
                     gate = c(0, 0, 1)),

  "CD16- Monocytes" = list(markers = c("CD14", "FCGR3A"),
                     gate = c(1, 0)),

  "CD16+ Monocytes" = list(markers = c("CD14", "FCGR3A"),
                     gate = c(1, 1)),

  "DC" = list(markers = c("CD3E", "CD14", "CD19", "NCAM1", "HLA-DRA"),
                     gate = c(0, 0, 0, 0, 1)),

  "Erythrocytes" = list(markers = c("HBA1"),
                     gate = c(7))
)
```

```r
classify <- function(dat, types) {
  class <- rep(NA, nrow(dat))
  res <- foreach(i = 1:length(types)) %do% {
    m <- types[[i]][["markers"]]
    cl <- foreach(j = 1:length(types[[i]][["markers"]]), .combine = "&") %do% {
      if(types[[i]][["gate"]][j]) {
        return(dat[ , m[j]] > types[[i]][["gate"]][j] - 1)
      } else {
        return(dat[ , m[j]] == 0)
      }
    }
    if(length(which(cl)) > 0) {
      class[which(cl & !is.na(class))] <- "AMBIG"
      class[which(cl & is.na(class))] <- names(types)[i]
    }
  }
  class
}

types <- classify(dat.imp, cellTypeDefs)
types[ which(types == "AMBIG") ] <- NA
types[ which(is.na(types)) ] <- "Unknown"
```

Note that the cell type definitions we have specified successfully assigns > 81% of cells to a single cell type, with less than 6% of cells assigned to more than one cell type (these abmiguous cell type assignments are removed). Some of the remaining ~17% of cells may be unassigned due to technical drop outs, while the others may be cell types we did not defined such as granulocytes that were not successfully removed by the ficoll gradient isolation of PBMCs, circulating epithelial cells, etc. Some may also represent empty droplets containing just background RNA from lysed cells that made it past our data filtering step.

For figures legends and other plots, we want the cell types to be in a specific order that is consistent and intuitive. So we define a function to set the order of the cell type labels for any given list of cell types. We also want the cell types to have consistent colors accross figures where applicable, so we create a helper function for that too.

```r
cellTypes <- c('B Cells',
               'CD4+ Naive T',
               'CD4+ Memory T',
               'CD4+ Effector T',
               'CD4+ Reg T',
               'CD8+ Naive T',
               'CD8+ Memory T',
               'CD8+ Effector T',
               "NKT CD4+",
               "NKT CD8+",
               "NKT CD4- CD8-",
               'NK',
               'CD16- Monocytes',
               'CD16+ Monocytes',
               'DC',
               'Erythrocytes',
               'Unknown')

fixOrder <- function(x) {
```

```r
  order <- cellTypes[which(cellTypes %in% unique(x))]
  factor(x, levels=order)
}

getCellPallette <- function(x) {
  x <- factor(x)
  cols<-c('#FFEE58',
          '#E65100',
          '#F57F15',
          '#FF8A65',
          '#FFCC80',

          '#0277BD',
          '#0097A7',
          '#00BCD4',

          '#1A237E',
          '#512DA8',
          '#9C27B0',
          '#9FA8DA',

          'gray', '#dddddd', "black", "red")

  ix <- match(levels(x), cellTypes)
  return(cols[ix])
}

types <- fixOrder(types)
saveRDS(types, "Final_Data/cell_types.rds")
```

To validate our scRNASeq analysis, we also performed FACS analysis on the same samples to directly measure the surface marker expression of major lymphocyte population markers. The percentages of the lymphocyte gate comprised of each subtype as determined by FACS are included in the metadata file. Proportion of lymphocytes in each population as assigned by FACS vs. scRNASeq is compared in Panel B.

```r
# load our data if we haven't already
load_init()

md <- readRDS("Final_Data/metadata_facs.rds")

options(scipen = 1e+06)
fig2B <- function() {
    # accumulate all our cell counts per sample, dropping
    # non-lymphocytes
    ids <- gsub("\\..*", "", rownames(dat.imp))
    sc.counts <- as.data.frame.matrix(table(ids, types))[, -c(16:17)]

    # aggregate some sub-populations to match flow populations
    sc.counts$`T Cells` <- sc.counts$`CD4+ Naive T` + sc.counts$`CD4+ Memory T` +
        sc.counts$`CD4+ Effector T` + sc.counts$`CD4+ Reg T` +
        sc.counts$`CD8+ Naive T` + sc.counts$`CD8+ Memory T` +
        sc.counts$`CD8+ Effector T` + sc.counts$`NKT CD4+` +
        sc.counts$`NKT CD8+` + sc.counts$`NKT CD4- CD8-`
```

```r
sc.counts$`CD4+ T Cells` <- sc.counts$`CD4+ Naive T` + sc.counts$`CD4+ Memory T` +
    sc.counts$`CD4+ Effector T` + sc.counts$`CD4+ Reg T` +
    sc.counts$`NKT CD4+`

sc.counts$`CD8+ T Cells` <- sc.counts$`CD8+ Naive T` + sc.counts$`CD8+ Memory T` +
    sc.counts$`CD8+ Effector T` + sc.counts$`NKT CD8+`

# and convert to % of lymphocytes
sc.lymphs <- table(grepl("CD4", types) | grepl("CD8", types) |
    grepl("B Cell", types) | grepl("NK", types), ids)[2,
    ]

sc.props <- sweep(sc.counts, 1, sc.lymphs, "/")
sc.props <- as.data.frame.matrix(sc.props)
sc.props <- sc.props[match(md$id, rownames(sc.props)), ]

# sanity check
all.equal(as.character(md$id), rownames(sc.props))

# extract the populations we are interested in
sc <- data.frame(`B Cells` = sc.props$`B Cells`)
flow <- data.frame(`B Cells` = md$`B Cells`)

# put space back in column names
colnames(sc)[1] <- "B Cells"
colnames(flow)[1] <- "B Cells"

sc$`T Cells` <- sc.props$`T Cells`
flow$`T Cells` <- md$`T Cells`

sc$`CD4+ T Cells` <- sc.props$`CD4+ T Cells`
flow$`CD4+ T Cells` <- md$`CD4+ T Cells`

sc$`CD8+ T Cells` <- sc.props$`CD8+ T Cells`
flow$`CD8+ T Cells` <- md$`CD8+ T Cells`

sc$`NK Cells` <- sc.props$NK
flow$`NK Cells` <- md$`NK Cells`

# assemble, melt, and plot
flow$ID <- md$id
sc$ID <- md$id

flow_m <- melt(flow, id.vars = c("ID"))
sc_m <- melt(sc, id.vars = c("ID"))

dat <- data.frame(ID = flow_m$ID, Population = flow_m$variable,
    Flow = flow_m$value, scRNASeq = sc_m$value)
ggplot(dat, aes(x = Flow, y = scRNASeq)) + geom_smooth(method = "lm",
    formula = y ~ x, color = "#cccccc", se = FALSE) + geom_point() +
    stat_cor(method = "pearson", aes(label = paste(..rr.label..,
        fp(..p..), sep = "~`,`~")), size = 3, label.x = 0,
        label.y = 0.95, digits = 3, color = "black") + facet_wrap(~Population,
```

```r
        scales = "free", ncol = 3) + scale_x_continuous(limits = c(0,
        1)) + scale_y_continuous(limits = c(0, 1)) + xlab(label = "Flow Cytometry") +
        theme_bw(base_size = 10) + my_theme + my_theme_wide_lab +
        theme(axis.text.x = element_text(angle = 315, hjust = 0),
            legend.position = "none")
}

F2B <- fig2B()
png("fig2_b.png", height = 1500, width = 2000, type = "cairo",
    res = 300)
print(F2B)
dev.off()
```

Next we wanted to visualize the clustering of cells in two dimensional space based on global gene expression in order to get a sense for whether we were capturing major biological signals in the the scRNASeq data. What we are hoping for is that there is minimal clustering by patient ID (after batch correction) and predominant clustering by cell type. We also see whether there are substantial clusters that distinguish cells from surviving vs. non-surving patients at this level of analysis (if not, deeper interrogation within cell types may be required).

```r
load_init()
types <- readRDS("Final_Data/cell_types.rds")
# types is the type of each cell, defined above
pal_cell <- getCellPallette(types)

# get palette of random colors for patient ID
colors = grDevices::colors()[grep("gr(a|e)y", grDevices::colors(),
    invert = T)]
set.seed(1010)
pal_id <- sample(colors, 38)

# and a two level palette for survival
pal_surv <- c("#1F70B0", "#BE1D2A")

umap <- readRDS("Final_Data/PBMC_merged_filtered_alra_pc_harmony_umap.rds")
ix <- which(types == "Unknown")
umap <- umap[-ix, ]
types <- types[-ix]
dat <- dat.imp[-ix, ]
ix.cell <- match(umap$id, md$Paper_ID)

a1 <- plotUmap(umap$UMAP1, umap$UMAP2, color = umap$id, label = "Normalized, color = id",
    palette = pal_id)
a2 <- plotUmap(umap$UMAP1, umap$UMAP2, color = types, label = "Normalized, color = cell type",
    palette = pal_cell, legend.position = "bottom")
a3 <- plotUmap(umap$UMAP1, umap$UMAP2, color = md$Survived[ix.cell],
    label = "Normalized, color = survival", palette = pal_surv,
    legend.position = "bottom")

# extract the legends to put them in their own panels
leg1 <- get_legend(a2 + guides(color = guide_legend(override.aes = list(size = 5,
    alpha = 1), ncol = 3, title = "Cell Type")) + theme(legend.text = element_text(margin = margin(l = [illegible]
    r = 10, unit = "pt")), legend.title = element_text(face = 2,
```

```
    margin = margin(r = 10, unit = "pt"))))

leg2 <- get_legend(a3 + guides(color = guide_legend(override.aes = list(size = 5,
    alpha = 1), title = "Survival")) + theme(legend.text = element_text(margin = margin(l = 2,
    r = 10, unit = "pt")), legend.title = element_text(face = 2,
    margin = margin(r = 10, unit = "pt"))))

a2 <- a2 + theme(legend.position = "none")
a3 <- a3 + theme(legend.position = "none")

# and assemble the plot

png("fig2_cde.png", height = 1500, width = 3000, type = "cairo",
    res = 300)
leg <- plot_grid(leg1, leg2, nrow = 1, rel_widths = c(0.7, 0.3))
p <- plot_grid(a1, a2, a3, nrow = 1, labels = c("", "", ""))
p <- plot_grid(p, leg, nrow = 2, rel_heights = c(0.7, 0.3))
print(p)
dev.off()
```

#### Figure 3

Figure 3, stratified by surviving vs. non-surviving patients (72 hours).

```
# load our data if we haven't already
load_init()

types <- readRDS("Final_Data/cell_types.rds")

fig3 <- function() {
    ids <- gsub("\\..*", "", rownames(dat.imp))

    # cell counts per patient, dropping unknown cells and
    # erythrocytes
    sc.counts <- as.data.frame.matrix(table(ids, types))[, -c(16:17)]

    # now convert to proportion of total cells
    totals <- apply(sc.counts, 1, sum)
    sc.counts.p <- sweep(sc.counts, 1, totals, "/")

    ix <- match(rownames(sc.counts.p), as.character(md$Paper_ID))
    # sanity check
    all.equal(rownames(sc.counts.p), as.character(md$Paper_ID)[ix])

    sc.counts.p$Survival <- md$Survived[ix]
    sc.counts.p$ID <- md$Paper_ID[ix]

    dat_m <- melt(sc.counts.p, id.vars = c("Survival", "ID"))
    dat_m$variable <- fixOrder(dat_m$variable)
```

```
    # now calculated p-value for t-test between survival groups
    # for each cell type, adjusting for multiple comparisons and
    # also format resulting labels and locations for plotting
    stat.test <- dat_m %>% group_by(variable) %>% t_test(value ~
        Survival)
    stat.test$p.adj <- p.adjust(method = "holm", stat.test$p)
    stat.test$yloc <- 0.75 * unlist(tapply(dat_m$value, INDEX = dat_m$variable,
        max))
    stat.test$x <- rep(1, nrow(stat.test))
    stat.test$p <- paste0("       p=", format(round(stat.test$p,
        3), nsmall = 3), "\n", "adj. p=", format(round(stat.test$p.adj,
        3), nsmall = 1))
    median2 <- function(x) {
        t <- median(x, na.rm = TRUE)
        return(data.frame(ymin = t, ymax = t, y = t))
    }

    # And plot.  Colors will match the colors in Figure 2, and
    # Figure 3A
    ggplot(dat_m, aes(x = Survival, y = value, color = variable)) +
        stat_summary(fun.data = median2, size = 0.5, geom = "crossbar",
            width = 0.6) + stat_pvalue_manual(stat.test, label = "p",
        remove.bracket = TRUE, x = "x", hjust = 0, y.position = "yloc",
        tip.length = 0, size = 3.5, color = "#444444") + geom_jitter(width = 0.07) +
        scale_color_manual(values = getCellPallette(dat_m$variable)) +
        ylab("Proportion of all Cells") + facet_wrap(~variable,
        scales = "free", nrow = 5) + my_theme + my_theme_wide_lab +
        theme(legend.position = "none")
}

F3 <- fig3()

png("fig3.png", height = 3000, width = 2000, type = "cairo",
    res = 300)
print(F3)
dev.off()
```

#### Figure 4

As none of the major cell types seemed predictive of survival, we wanted to drill into each cell type and find biological processes and surface markers that might distinguish surviving vs. non-surviving patients. Panel B displays the results of this analysis. For identification of differentially expressed genes and associated biological processes, we limited our analysis to those genes that had variable expression defined as having an expression level normalized robust z-score greater than 2. This approach is the same as that described in Zheng et al. (2017) and Macosko, et al. (2015).

Here is the function we used to calculate the dispersion for each gene:

```
dispersion <- function(x, dim = 2, verbose = FALSE) {

    if (verbose)
        message("Calculating means (1 of 4)")
    means <- apply(x, dim, mean, na.rm = TRUE)
```

```r
    if (verbose)
        message("Calculating dispersion (2 of 4)")
    disp <- apply(x, dim, function(x) {
        var(x, na.rm = TRUE)/mean(x, na.rm = TRUE)
    })

    if (verbose)
        message("Binning (3 of 4)")
    bins <- cut(means, quantile(means, seq(0, 1, len = 20)),
        include.lowest = TRUE)

    if (verbose)
        message("Normalizing (4 of 4)")
    rv <- tapply(disp, bins, function(y) {
        abs(y - median(y))/mad(y)
    })
    rv <- unlist(rv)
    names(rv) <- gsub("\\[.*\\]\\.", "", names(rv))
    rv
}
```

We then used this function to identify the variable gene set based on cells of known type (as defined above) that were not erythrocytes.

```r
# load our data if we haven't already
load_init()

types <- readRDS("Final_Data/cell_types.rds")

ix <- which(!types %in% c("Erythrocytes", "Unknown"))
dat <- dat.imp[ix, ]
disp <- dispersion(dat)
ix <- which(disp > 2)
gg <- gsub(".*\\]\\.", "", names(disp))
gg <- gg[ix]

# remove pseudogenes and non-coding genes from a list of
# genes
expGenes <- function(x) {
    x <- x[-which(grepl("A\\w\\d*\\.\\d", x))]
    x <- x[-which(grepl("C\\d{1,2}orf", x))]
    x <- x[-which(grepl("^LINC\\d+$", x))]
    x
}
gg <- expGenes(gg)
ix <- which(colnames(dat) %in% gg)
dat <- dat[, ix]
saveRDS(dat, file = "Final_Data/PBMC_merged_filtered_alra_variable_genes.rds")
```

Now we need helper functions to calculate the proportion of cells within each subtype that express each gene for each patient.

```r
# load our data if we haven't already
load_init()

applyById <- function(x, ids, FUN = function(x) {
    sum(x > 0)
}) {
    ids.x <- gsub("\\..*", "", names(x))
    rv <- foreach(i = ids, .combine = c) %do% {
        ix <- which(ids.x == i)
        if (length(ix)) {
            FUN(x[ix])
        } else {
            0
        }
    }
    names(rv) <- ids
    rv
}

# convenience wrapper
countsById <- function(x, ids) {
    applyById(x, ids, length)
}

# convenience alias
positiveById <- function(x, ids) {
    applyById(x, ids)
}

R_na_zero <- function(x) {
    ix <- which(is.na(x))
    if (length(ix))
        x[ix] <- 0
    x
}

genesProp <- function(types, dat, cluster = TRUE) {
    calcRatio <- function(x, ix.s, ix.d, ids.s, ids.d) {
        pos.s <- positiveById(x[ix.s], ids.s)
        tot.s <- countsById(x[ix.s], ids.s)
        # if there are no cells for an id, assum proportion is zero
        prop.s <- R_na_zero(pos.s/tot.s)

        pos.d <- positiveById(x[ix.d], ids.d)
        tot.d <- countsById(x[ix.d], ids.d)
        prop.d <- R_na_zero(pos.d/tot.d)

        prop.tot <- sum(pos.s, pos.d)/sum(tot.s, tot.d)

        ratio = log2(median(prop.d)/median(prop.s))
        if (is.nan(ratio))
            ratio <- 0
```

```
        data.frame(logratio = ratio, Survived = median(prop.s),
            Died = median(prop.d), prop.positive = prop.tot,
            p = suppressWarnings(wilcox.test(prop.s, prop.d)$p.value))
    }

    ids <- gsub("\\..*", "", rownames(dat))
    ix <- match(ids, md$Paper_ID)
    survival <- md$Survived[ix]

    if (cluster) {
        cl <- makeCluster(detectCores() - 1)
        registerDoParallel(cl)
    }

    ratios <- foreach(t = levels(types), .packages = c("dplyr",
        "foreach"), .export = c("applyById", "positiveById",
        "countsById", "R_na_zero")) %dopar% {
        ix.type <- which(types == t)
        dat.type <- dat[ix.type, ]
        ix.s <- which(survival[ix.type] == "Survived")
        ix.d <- which(survival[ix.type] == "Died")
        ids.s <- unique(ids[which(survival == "Survived")])
        ids.d <- unique(ids[which(survival == "Died")])
        rv <- apply(dat.type, 2, calcRatio, ix.s, ix.d, ids.s,
            ids.d)
        rv <- do.call(rbind, rv)
        rv$p.adj <- p.adjust(rv$p, "BH")
        rv$gene <- colnames(dat)
        rv
    }
    if (cluster) {
        stopCluster(cl)
    }
    names(ratios) <- levels(types)
    ratios
}

types <- readRDS("Final_Data/cell_types.rds")
types.p <- factor(types[which(!types %in% c("Erythrocytes", "Unknown"))])
dat <- readRDS("Final_Data/PBMC_merged_filtered_alra_variable_genes.rds")
props <- genesProp(types.p, dat)
names(props) <- levels(types.p)
saveRDS(props, file = "Final_Data/gene_props.rds")
```

We can then plot a heatmap of the gene proportions. Red genes are those expressed in a higher proportion of cells from deceased patients, while blue genes are those expressed in a higher proportion of cells from surviving patients. The intensity of the color is proportional to the adjusted (FDR) p-value (t-test) for each gene (within each subtype).

```
fig4_a_hm <- function() {
    props <- readRDS("Final_Data/gene_props.rds")
    # take 1 - adjusted pvalue (so the most significant genes are
    # the extremes of the scale), and set sign to the direction
```

```
    # of the fold change.
    pv <- foreach(d = props, .combine = cbind) %do% {
        (1 - d$p.adj) * sign(d$logratio)
    }

    pv[which(is.na(pv))] <- 0
    colnames(pv) <- levels(types.p)
    rownames(pv) <- props$`B Cells`$gene

    # remove completely uninformative genes
    ix.rm <- which(apply(pv, 1, sum) == 0)
    pv <- pv[-ix.rm, ]

    hm <- pheatmap(pv, silent = TRUE, legend = FALSE, border_color = NA,
        cluster_rows = TRUE, cluster_cols = FALSE, show_rownames = FALSE,
        color = colorRampPalette(c("#005AC8", "white", "#FA2800"))(9),
        treeheight_col = 10, treeheight_row = 30)

    # save the dendrogram for further analysis below
    dg <- rev(as.dendrogram(hm$tree_row))
    saveRDS(dg, file = "Final_Data/gene_prop_dendrogram.rds")

    # grab the legend
    hm.leg <- pheatmap(pv, silent = TRUE, breaks = c(seq(-1,
        1, 0.2)), legend_breaks = seq(-1, 1, 0.2), legend_labels = format(seq(-1,
        1, 0.2), nsmall = 2), legend = TRUE, color = colorRampPalette(c("#005AC8",
        "white", "#FA2800"))(9))$gtable$grobs[[6]]
    # hm.leg <- as.ggplot(hm.leg) + theme(plot.margin =
    # unit(c(1,0,0,0), 'cm'))
    hm.g <- as.ggplot(hm)
    plot_grid(hm.g + theme(plot.margin = unit(c(1, -0.2, 0, 1),
        "cm")), grobTree(hm.leg, vp = viewport(x = 0.7, y = 1.75,
        angle = 90)), ncol = 1, nrow = 2, labels = c(" ", " "),
        label_size = 10, rel_heights = c(1, 0.15))
}

F4A_pre <- fig4_a_hm()
```

In an effort to functionalize the gene clusters, we performed Gene Ontology term enrichment analysis on the genes in each of the major subclusters. We create a helper function that accepts a dendrogram, and the index of a subtree, and then runs GO enrichment analysis on the genes in that subtree. The function also optionally plots the subtree so you can visually verify that you know which subtree is being analyzed by cross referencing to the whole dendrogram.

```
library(dendextend)
library(clusterProfiler)
library(org.Hs.eg.db)
library(stringr)

# get 1:1 mapping of gene symbol to entrez gene, dropping NAs
symToEG <- function(x) {
    x <- x[which(x %in% ls(org.Hs.egSYMBOL2EG))]
    x <- mget(x, org.Hs.egSYMBOL2EG)
```

```r
    unlist(lapply(x, function(i) {
        i[1]
    }))
}

# k is the number of subtrees to cut the dendrogram into, i
# is the subtree of interest. If heatmap=TRUE, exp must be
# supplied (the full data matrix originally used for
# clustering)
dendGo <- function(dend, k, i, heatmap = FALSE, go = TRUE, exp = NULL) {
    gt <- NULL
    clusters <- dendextend::cutree(dend, k)
    genes <- labels(dend)
    labels(dend) <- clusters[order.dendrogram(dend)]
    ix <- which(clusters[order.dendrogram(dend)] == i)
    labels(dend) <- genes
    gg <- labels(dend)[ix]
    if (heatmap) {
        if (is.null(exp)) {
            stop("If heatmap=TRUE, must supply data matrix exp")
        }
        p <- pheatmap(exp[which(rownames(exp) %in% gg), ], cluster_cols = FALSE,
            border_color = NA, cluster_rows = TRUE, show_rownames = FALSE,
            color = colorRampPalette(c("#005AC8", "white", "#FA2800"))(11),
            treeheight_col = 10, treeheight_row = 50)
        print(p)

    }
    if (go) {
        go_test <- enrichGO(symToEG(gg), OrgDb = "org.Hs.eg.db",
            pvalueCutoff = 0.05)
        go_test@result$Description <- go_test@result$Description
        gt <- go_test
        # return(dotplot(go_test, font.size=10) +
        # scale_size_continuous(range = c(1, 6)))
    }
    return(gt)
}
```

We then step through each of 12 (mutually exclusive, but exhaustive) subtrees and test for enriched GO terms. Although the number of subtrees is arbitrary, the identification of subtrees once the number is chosen is systematic and automated, performed using Tal Galili's `dendextend` package.

```r
library(dendextend)
library(doParallel)

dg <- readRDS(file = "Final_Data/gene_prop_dendrogram.rds")
cl <- makeCluster(detectCores() - 1)
registerDoParallel(cl)

go_enrich <- foreach(i = 1:12, .packages = c("dendextend", "clusterProfiler",
    "stringr")) %dopar% {
    dendGo(dg, 12, i)
```

```r
}

stopCluster(cl)
saveRDS(go_enrich, "Final_Data/go_enrichment.rds")
```

It would be helpful to create an annotation bar that will indicate where exactly the sub trees are, and to label these bars with any significantly enriched GO terms. Because space is limited, a maximum of 2 terms are annotated on the plot, and terms are truncated for length as needed.

```r
topTerms <- function(enr, n = 3, strlen = 50) {
    termlist <- foreach(i = 1:length(enr)) %do% {
        r <- enr[[i]]@result

        # truncate and remove duplicate terms that are identical
        # after truncation
        r$Description <- str_trunc(r$Description, strlen)
        ix.rm <- which(duplicated(r$Description))
        r <- r[-ix.rm, ]
        ix <- which(r$p.adjust < 0.05)
        if (length(ix)) {
            if (length(ix) > n) {
                terms <- r$Description[ix][1:n]
            } else {
                terms <- r$Description[ix]
            }
        } else {
            terms <- ""
        }
        terms
    }
    termlist
}

dendnote <- function(dend, k, offset = 0.125, cols = c("gray",
    "black"), reverse = TRUE, sort.labels = FALSE, enr = NULL) {
    if (reverse) {
        dend <- rev(dend)
    }
    order <- order.dendrogram(dend)
    clusters <- cutree(dend, k)
    levels <- unique(clusters[order])
    dn <- data.frame(labels = unique(clusters[order]))
    if (length(enr)) {
        dn$label = sapply(topTerms(enr, 2)[levels], paste, collapse = "\n")
    } else {
        dn$label = ""
    }
    if (sort.labels) {
        dn$id <- sort(levels, decreasing = TRUE)
    } else {
        dn$id <- levels
    }
    dn$at <- which(!duplicated(clusters[order]))
```

```
    dn$length <- as.vector(table(clusters[order])[levels])
    dn$at_ann <- dn$at + dn$length
    dn$at <- dn$at + 0.5 * dn$length
    dn$col <- cols
    dn$offset <- c(0, offset)
    ggplot(dn, aes(x = 1 + offset, y = at, height = length, fill = col,
        label = id)) + geom_tile(width = 0.25) + geom_text(aes(x = 1.8,
        col = col), size = 4.5) + geom_text(aes(x = 2.5, y = at,
        label = label, col = col), lineheight = 0.8, size = 4,
        hjust = 0, vjust = 0.5) + scale_fill_manual(values = cols) +
        scale_color_manual(values = cols) + coord_cartesian(xlim = c(0,
        8), clip = "off") + theme(axis.line = element_blank(),
        axis.text.x = element_blank(), axis.text.y = element_blank(),
        axis.ticks = element_blank(), axis.title.x = element_blank(),
        axis.title.y = element_blank(), legend.position = "none",
        panel.background = element_blank(), panel.border = element_blank(),
        panel.grid.major = element_blank(), panel.grid.minor = element_blank(),
        plot.background = element_blank())

}
```

We can then add the annotation bar to our heatmap.

```
dg <- readRDS(file = "Final_Data/gene_prop_dendrogram.rds")
go_enrich <- readRDS("Final_Data/go_enrichment.rds")

# fig4b panel 1 with GO annotation
fig4a_ann <- function() {
    blank <- grid.rect(gp = gpar(col = "white"))
    dend <- plot_grid(dendnote(dg, 12, reverse = FALSE, sort.labels = TRUE,
        enr = go_enrich, cols = c("black", "#006666")) + theme(plot.margin = margin(0.25,
        2, 2.3, -0.8, "cm")), blank, nrow = 2, rel_heights = c(1,
        0.15))

    plot_grid(F4A_pre, dend, rel_widths = c(1.2, 1))
}
F4A <- fig4a_ann()
```

We did not have room in the figure to display every significant GO term. But of course, we want to report them, and we do so in supplemental table S4. For figure 4 in the manuscript, we wanted the node ids to be in numerical order, top to bottom. But this is not the way `cutree` numbers the subtrees (it numbers them hierachically as it encounters them). Below we create a table of significant GO terms with both the original node id as assigned by `cutree` and the node id as presented in the figure.

```
go_enrich <- readRDS("Final_Data/go_enrichment.rds")
dg <- readRDS(file = "Final_Data/gene_prop_dendrogram.rds")
clusters <- cutree(dg, 12)
order <- order.dendrogram(dg)
pub_id <- match(c(1:12), rev(unique(clusters[order])))

res <- NULL
for (i in 1:length(go_enrich)) {
    r <- go_enrich[[i]]@result
```

```
    ix <- which(r$p.adjust < 0.05)
    if (length(ix)) {
        res <- rbind(res, cbind(pub_id[i], i, r$Description[ix],
            r$p.adjust[ix]))
    }
}
colnames(res) <- c("Node_ID_pub", "Node_ID_orig", "GO_Term",
    "adj_p")
write.table(res, quote = FALSE, row.names = FALSE, sep = "\t",
    file = "Final_Data/table_s4.tab")
```

We also wanted to examine cytokines specifically. Cytokines were identified manually based on published lists and our own experience. First we extract these cytokines from the gene expression matrix, again calculate proportions of positive cells within each cell type, and generate another heatmap. Note that the heatmap was split into two panels (predominantly upregulated in Survived, and predominatnyl upregulated in Died) using image editing software.

```
types <- readRDS("Final_Data/cell_types.rds")
ix <- which(!types %in% c("Erythrocytes", "Unknown"))
types.p <- factor(types[ix])

gg.cyt <- unique(readLines("Final_Data/cytokine_list.txt"))
dat <- dat.imp[ix, gg.cyt]
props <- genesProp(types.p, dat)
names(props) <- levels(types.p)
saveRDS(props, file = "Final_Data/gene_props_cytokines.rds")
```

We can then plot a heatmap of these genes. As it happens, they split nicely into two groups.

```
props <- readRDS("Final_Data/gene_props_cytokines.rds")

fig4b <- function() {
    props <- readRDS(file = "Final_Data/gene_props_cytokines.rds")
    # take 1 - adjusted pvalue (so the most significant genes are
    # the extremes of the scale), and set sign to the direction
    # of the fold change.
    pv <- foreach(d = props, .combine = cbind) %do% {
        (1 - d$p.adj) * sign(d$logratio)
    }

    pv[which(is.na(pv))] <- 0
    colnames(pv) <- levels(types.p)
    rownames(pv) <- props$`B Cells`$gene

    breaks <- seq(-1, 1, 0.19)
    cols <- colorRampPalette(c("#005AC8", "white", "#FA2800"))(length(breaks))

    hm <- pheatmap(pv, silent = TRUE, legend = FALSE, border_color = NA,
        cluster_rows = TRUE, cluster_cols = FALSE, show_rownames = TRUE,
        fontsize_row = 8, color = colorRampPalette(c("#005AC8",
            "white", "#FA2800"))(11), treeheight_col = 10, treeheight_row = 30)

    dg <- rev(as.dendrogram(hm$tree_row))
```

```
    saveRDS(dg, file = "Final_Data/gene_prop_dendrogram.rds")
    clusters <- cutree(dg, 2)
    ix1 <- which(clusters == 1)
    ix2 <- which(clusters == 2)

    hm1 <- pheatmap(pv[ix1, ], silent = TRUE, legend = FALSE,
        border_color = NA, cluster_rows = TRUE, cluster_cols = FALSE,
        show_rownames = TRUE, fontsize_row = 8, color = cols,
        breaks = breaks, treeheight_col = 10, treeheight_row = 30)
    hm2 <- pheatmap(pv[ix2, ], silent = TRUE, legend = FALSE,
        border_color = NA, cluster_rows = TRUE, cluster_cols = FALSE,
        show_rownames = TRUE, fontsize_row = 8, color = cols,
        breaks = breaks, treeheight_col = 10, treeheight_row = 30)

    hm1 <- as.ggplot(hm1)
    hm2 <- as.ggplot(hm2)
    plot_grid(hm2 + theme(plot.margin = unit(c(1, 0, 0, 0.5),
        "lines")), hm1 + theme(plot.margin = unit(c(1, 0, 1.8,
        0.5), "lines")), ncol = 2, hjust = -0.25, label_size = 10,
        labels = c("Predominantly Up in Survived", "Predominantly Up in Died"))
}

F4B <- fig4b()
```

And we can assemble the figure. Note that some additional annotation to the legend was added after the fact for the final figure in the paper.

```
png("fig4.png", width = 4500, height = 3000, res = 300)
plot_grid(F4A, F4B + theme(plot.margin = unit(c(1, 0, 1.8, 0.5),
    "lines")), ncol = 2, rel_widths = c(0.45, 0.55), labels = c("A",
    "B"))
dev.off()
```

#### Figure 5

In order to facilitate future isolation of interesting cell subpopulations by FACS, we would also like to focus in on surface markers specifically. We obtained a list of known clusters of differentiations, and their alternative gene symbols (if any) from the Cell Surface Protein Atlas and used this to filter our gene list. The resulting list of genes is presented as Supplemental Table S5 in our manuscript, and is also available as `Final_Data/table_s5.txt` in the github repository.

We are most interested in those surface markers that exhibited a strong fold in proportion of positive cells between surviving and non-surviving patients and that have a p-value corresponding to a relatively low false discovery rate. For our purposes, we will screen for surface markers with a fold change of at least 1.5 (in either direction) and p-value $< 0.01$. The corresponding candidate markers are shown in figure 5A. Note that we limit our analysis to those surface markers that are among the highly variable genes defined earlier.

```
types <- readRDS("Final_Data/cell_types.rds")
ix <- which(!types %in% c("Erythrocytes", "Unknown"))
types.p <- factor(types[ix])

dat <- readRDS("Final_Data/PBMC_merged_filtered_alra_variable_genes.rds")
```

```
cds <- read.delim("Final_Data/table_s5.txt", as.is = TRUE, header = TRUE)

ix.cd <- which(colnames(dat) %in% cds$Gene_Symbol | grepl("^CD\\d",
    colnames(dat)))

props_cd <- genesProp(types.p, dat[, ix.cd])
names(props_cd) <- levels(types.p)
saveRDS(props_cd, file = "Final_Data/cd_props.rds")
```

We can then plot the enriched surface markers, sizing the points by the proportion of all cells for each cell type that express each marker so that we can visualize which are the major populations.

```
props_cd <- readRDS("Final_Data/cd_props.rds")
tt <- foreach(l = props_cd, .combine = cbind) %do% {
    l$p
}

# remove completely uninformative markers
nx <- apply(tt, 1, function(x) {
    sum(x < 0.01, na.rm = TRUE)
})
ix <- which(nx > 0)
dat <- lapply(props_cd, function(x) {
    x[ix, ]
})

# format the data for plotting
dat.m <- melt(dat, id.vars = colnames(dat[[1]]))
ix <- which(abs(dat.m$logratio) < 0.4)
dat.m <- dat.m[-ix, ]
dat.m <- dat.m[order(dat.m$p, decreasing = TRUE), ]
dat.m$gene <- factor(dat.m$gene, levels = unique(dat.m$gene))
dat.m$p <- sign(-dat.m$logratio) * -log(dat.m$p)

F5A <- ggplot(dat.m, aes(x = L1, y = gene, size = prop.positive,
    color = p)) + geom_point() + scale_colour_gradientn(colours = c("#C02222",
    "white", "#176BA0")) + xlab("Cell Type") + ylab("Surface Marker") +
    theme_bw() + my_theme + theme(axis.text.x = element_text(angle = 270,
    hjust = 0, vjust = 0.5)) + guides(color = guide_colorbar(title = "-log(p-value)",
    reverse = TRUE), size = guide_legend(title = "Proportion Positive"))
```

We would like to perform survival analysis based on the novel markers we have selected. Here we create a helper function to perform the analysis and plot the results.

```
library(survival)

survAnalysis <- function(cellp, cutoff = NULL, days = 3, pop.name = "Positive cells / total lymphs:") {
    md <- readRDS("Final_Data/metadata_clin_cyt.rds")
    md <- md[which(md$Paper_ID %in% names(cellp)), ]
    ix <- match(md$Paper_ID, names(cellp))
    md$cellp <- cellp[ix]
    md$surv_time[which(md$surv_time > (days * 24))] <- days *
        24
```

```
    event <- factor(md$surv_time < days * 24, labels = c("Survived",
        "Died"))
    surv <- Surv(time = md$surv_time, event = event == "Died")

    if (is.null(cutoff)) {
        cutoff <- median(cellp)
    }
    md$surv <- surv
    md$high_count <- factor(md$cellp > cutoff, labels = c("High",
        "Low"))
    fit <- coxph(surv ~ high_count, data = md)
    print(summary(fit))
    print(survdiff(surv ~ high_count, data = md))
    ggsurvplot(survfit(surv ~ high_count, data = md), color = "black",
        ggtheme = my_theme_no_space + my_theme_wide_lab + theme(legend.title = element_text(size = 10,
            face = "bold"), legend.text = element_text(size = 10)),
        data = md, pval = TRUE, pval.coord = c(0.6 * days * 24,
            0.1), pval.size = 3.5, linetype = "strata", risk.table = FALSE,
        legend.title = pop.name, legend.labs = c("Low", "High")) +
        xlab("Time (Hours)") + ylab("Survival")
}
```

We can then perform survival analysis based on CD52 expression in CD8+ NKT Cells.

```
load_init()

fig5b <- function() {
    dat <- readRDS("Final_Data/PBMC_merged_filtered_alra_variable_genes.rds")
    types <- readRDS("Final_Data/cell_types.rds")
    types.p <- factor(types[which(!types %in% c("Erythrocytes",
        "Unknown"))])
    ids <- gsub("\\..*", "", rownames(dat))
    pd <- data.frame(ids = ids, type = types.p)

    md <- readRDS("Final_Data/metadata_clin_cyt.rds")
    md <- md[which(md$group == "Initial"), ]
    ix <- match(pd$ids, md$Paper_ID)
    pd$Survival <- md$Survived[ix]
    pd$CD52 <- dat[, "CD52"] > 0

    cd8_nkt <- table(pd$id, pd$type)[, "NKT CD8+"]
    cd8_nkt_cd52 <- table(pd$id, pd$CD52, pd$type)[, 2, "NKT CD8+"]
    cellp <<- cd8_nkt_cd52/cd8_nkt
    p1 <- survAnalysis(cellp, pop.name = "CD8+ NKT/CD52+")[[1]]
    p1
}

F5B <- fig5b()
```

Finally, it would be interesting to see whether the proportion of CD8+ NKT cells that are CD52+, in addition to correlating with survival, also correlates with other key markers from our prior analysis includiing IL6 and Arterial pH. This is shown in panels C and D of Figure 5.

```r
load_init()

p1 <- ggplot(md, aes(x = cd52_prop, y = IL_6_19 + 1)) + geom_point() +
    stat_smooth(method = "lm") + ylab(bquote(log[10] ~ bold("IL-6") ~
    scriptstyle("(pg/ml)"))) + annotate("text", size = 3, fontface = 2,
    label = "p=0.011, r = -0.416", x = -Inf, y = Inf, hjust = -0.1,
    vjust = 1.5) + scale_y_continuous(minor_breaks = scales::extended_breaks(15),
    trans = "log10", breaks = c(0, 1, 10, 100, 1000, 10000, 1e+05),
    labels = function(x) {
        format(round(log10(x)), scientific = FALSE)
    }, expand = expand_scale(mult = c(0.1, 0.2), add = 0)) +
    xlab("CD8+ NKT/CD52+") + theme_bw() + my_theme + my_theme_pretty_grid +
    theme(plot.margin = unit(c(1, 1, 0.2, 1), "lines"))

p2 <- ggplot(md, aes(x = cd52_prop, y = ph_art)) + geom_point() +
    stat_smooth(method = "lm") + ylab(bquote("Arterial pH")) +
    annotate("text", size = 3, fontface = 2, label = "p=0.014, r = 0.396",
        x = -Inf, y = Inf, hjust = -0.1, vjust = 1.5) + scale_y_continuous(expand = expand_scale(mult =
    0.2), add = 0)) + xlab("CD8+ NKT/CD52+") + theme_bw() + my_theme +
    my_theme_wide_lab + my_theme_pretty_grid + theme(plot.margin = unit(c(1,
    1, 0.2, 1), "lines"))

F5CD <- plot_grid(p1, p2, ncol = 2, nrow = 1, align = "v", labels = c("C",
    "D"))
```

And we can assemble the complete figure…

```r
png("fig5.png", width = 1700, height = 3500, res = 300)
plot_grid(F5A, F5B + theme(plot.margin = unit(c(0.2, 1, 1, 1),
    "lines")), F5CD, nrow = 3, rel_heights = c(0.5, 0.3, 0.2),
    labels = c("A", "B", ""))
dev.off()
```

#### Figure 6

Figure 6 summarizes our analysis of a second cohort of patients assessed to validate our findings with respect to CD8+/CD52+ NKT cells.

Panel A is comprised of representative density plots illustrating the population shift between surviving and non-surviving patients.

```r
F6A <- ggplot() + theme_void()
```

Panel B again presents clinical data, but now stratified by proportion of CD8+ NKT cells that are also CD52. We used the same cutoff defined in the initial phase of the study to stratify the patients.

```r
md <- readRDS("Final_Data/metadata_clin_cyt.rds")
ix.val <- which(md$group == "Validation" & md$mode == "VA")
md <- md[ix.val, ]
md$High_CD52 <- md$cd52_prop > 0.7646
```

```
dat <- md[, c("age", "ph_art", "sofa", "mdrd_egfr_calc", "High_CD52",
    "surv_time")]
colnames(dat) <- c("Age", "Arterial pH", "SOFA", "eGFR", "CD52",
    "surv_time")
dat$CD52 <- factor(md$High_CD52, labels = c("Low CD52+", "High CD52+"))

fp <- function(p) {
    if (p < 0.001)
        return("p < 0.001")
    return(paste("p = ", round(p, 3)))
}

pvv <- foreach(i = 1:4) %do% {
    t.test(dat[, i] ~ dat$CD52)$p.value
}

pl <- foreach(i = 1:4) %do% {
    gd <- data.frame(CD52 = dat$CD52, y = dat[, i])
    ggplot(gd, aes(x = CD52, y = y)) + geom_jitter(width = 0.1,
        height = 0.1) + stat_summary(fun.y = median, color = "red",
        geom = "point", aes(group = 1), size = 3, show.legend = FALSE) +
        ylab(colnames(dat)[i]) + annotate("text", size = 3, fontface = 2,
        label = fp(as.numeric(pvv[i])), x = -Inf, y = Inf, hjust = -0.1,
        vjust = 1.5) + scale_y_continuous(expand = expand_scale(mult = c(0.1,
        0.2), add = 0)) + xlab("") + theme_bw() + my_theme +
        my_theme_wide_lab + my_theme_pretty_grid + theme(axis.text.x = element_text(angle = 315,
        hjust = 0.1, vjust = 0.5)) + theme(plot.margin = unit(c(0.1,
        1, 0.2, 0.2), "lines"))
}

F6B <- plot_grid(plotlist = pl, ncol = 2, nrow = 2, align = "v")
```

Panel C is survival analysis of the stratified patients.

```
md$surv_time_72 <- md$surv_time
md$surv_time_72[which(md$surv_time > 72)] <- 72
md$surv_72 <- Surv(md$surv_time_72, event = md$surv_time_72 <
    72)
print(survdiff(surv_72 ~ High_CD52, data = md))
print(summary(coxph(surv_72 ~ High_CD52, data = md)))
F6C <- ggsurvplot(survfit(surv_72 ~ High_CD52, data = md), color = "black",
    ggtheme = my_theme_no_space + my_theme_wider_lab + theme(legend.title = element_text(size = 10,
        face = "bold"), legend.text = element_text(size = 10)),
    data = md, pval = TRUE, pval.coord = c(0.6 * 3, 0.1), pval.size = 3.5,
    linetype = "strata", risk.table = FALSE, legend.title = "CD8+ NKT/CD52+",
    legend.labs = c("Low", "High")) + xlab("Time (Hours)") +
    ylab("Survival")

F6C <- F6C[[1]]
```

And we can assemble the plot:

```r
png("fig6.png", width = 1500, height = 3500, res = 300)
plot_grid(F6A, F6B + theme(plot.margin = unit(c(1, 1, 0, 1),
    "lines")), F6C + theme(plot.margin = unit(c(0.2, 1, 0, 1),
    "lines")), nrow = 3, rel_heights = c(0.33, 0.4, 0.27), labels = c("A",
    "B", "C"))
dev.off()
```
