## Supplemental tables and figures for "The inflammasome of circulatory collapse: single cell analysis of survival on extracorporeal life support"

**This PDF file includes:**

Tables S1 to S5

Figures S1, S2

**The Following Supplemental material are provided as separate Files:**

**Supplemental Data D2:** List of variable genes, provide in file Data_S2.tab

**Supplemental Data D3:** Details of data analysis is provided at final_analysis.html. The source file and supporting data are also provided online at https://github.com/vanandelinstitute/va_ecls.

| Cytokine | Median Died | Median Survived | Mean Died | Mean Survived | Wilcox Test (adj. p) | > 0 |
|---|---|---|---|---|---|---|
| IL-1β | 4.1 | 1.3 | 7.6 | 1.9 | 0.016 | 36 |
| IL-2 | 0.0 | 0.0 | 6.2 | 1.2 | 1.000 | 9 |
| IL-4 | 2.6 | 0.4 | 4.2 | 0.8 | 0.011 | 33 |
| IL-5 | 1.7 | 0.4 | 2.5 | 1.3 | 1.000 | 20 |
| IL-6 | 1337.3 | 149.7 | 19704.0 | 334.8 | 0.001 | 36 |
| IL-7 | 3.6 | 3.3 | 5.0 | 4.8 | 1.000 | 34 |
| IL-8 | 111.3 | 16.6 | 185.6 | 44.6 | 0.116 | 36 |
| IL-10 | 114.3 | 35.5 | 193.2 | 121.2 | 0.249 | 35 |
| IL-12 | 9.6 | 3.9 | 11.7 | 5.3 | 0.008 | 35 |
| IL-13 | 1.9 | 0.2 | 4.6 | 1.3 | 0.094 | 24 |
| IL-17 | 59.1 | 31.0 | 68.5 | 44.5 | 0.576 | 35 |
| G-CSF | 263.0 | 74.7 | 6014.0 | 539.8 | 0.116 | 36 |
| GM-CSF | 53.9 | 5.7 | 85.4 | 34.1 | 0.062 | 27 |
| IFNγ | 91.9 | 4.8 | 126.3 | 27.2 | 0.094 | 27 |
| MCP/MCAF | 287.9 | 114.0 | 511.0 | 152.3 | 0.058 | 34 |
| MIP-1β | 92.0 | 62.6 | 150.9 | 108.8 | 0.880 | 36 |
| TNFα | 47.2 | 9.1 | 107.0 | 22.5 | 0.011 | 36 |

**Table S1.** Plasma cytokine levels stratified by 72 hour survival. P-values are from the Wilcoxon rank-sum test, adjusted for multiple comparison using the method of Holm. The “>0” column indicated the number of patients (out of 36) for whom each cytokine was detectable. Levels below the detection limit of the assay were set to “0” for statistical analyses.

| | CD19 | CD3 | CD4 | CD8 | CD2 | CD57 | CD25 | FOXP3 | CD56 | CD14 | CD16 | HLA-DRA | HBA1 | |
|---|---|---|---|---|---|---|---|---|---|---|---|---|---|---|
| **B Cells** | + | - | | | | | | | | | | | | |
| **CD4 Naïve T** | | + | + | - | - | | +/-* | +/-* | - | | | | | * Cannot be CD25/FOXP3 double positive |
| **CD4 Memory T** | | + | + | - | + | - | +/-* | +/-* | - | | | | | * Cannot be CD25/FOXP3 double positive |
| **CD4 Effector T** | | + | + | - | + | + | +/-* | +/-* | - | | | | | * Cannot be CD25/FOXP3 double positive |
| **CD4 Regulatory T** | | + | + | - | | | + | + | - | | | | | |
| **CD8 Naïve T** | | + | - | + | - | | | | - | | | | | |
| **CD8 Memory T** | | + | - | + | + | - | | | - | | | | | |
| **CD8 Effector T** | | + | - | + | + | + | | | - | | | | | |
| **CD4+ NKT** | - | + | + | - | | | | | + | | | | | |
| **CD8+ NKT** | - | + | - | + | | | | | + | | | | | |
| **CD4- CD8- NKT** | - | + | - | - | | | | | + | | | | | |
| **Natural Killer** | - | - | | | | | | | + | | | | | |
| **Monocyte** | | | | | | | | | | + | +/- | | | |
| **Dendritic Cell** | - | - | | | | | | | - | - | | + | | |
| **Erythrocyte** | | | | | | | | | | | | | ++ | |

**Table S2.**
Classification scheme for PBMC subsets based on scRNASeq expression

**Table S3.**

Impact of imputation on slope (beta) and $R^2$ of correlation between FACS and scRNASeq based assignment of cells to major lymphocyte classes, as a proportion of all lymphocytes. P-value listed is for t-test for difference between either Beta or $R^2$ pre- and post-imputation.

| | **Pre-imputation** | | **Post-imputation** | |
|---|---|---|---|---|
| | **Beta** | **$R^2$** | **Beta** | **$R^2$** |
| **B Cells** | 0.95578 | 0.57140 | 0.82673 | 0.61830 |
| **T Cells** | 0.95161 | 0.75850 | 0.86627 | 0.70840 |
| **CD4+** | 0.56471 | 0.61950 | 0.82542 | 0.70800 |
| **CD8+** | 0.97919 | 0.41240 | 0.71510 | 0.36410 |
| **NK** | 0.90466 | 0.57290 | 1.12664 | 0.56950 |
| **Average** | 0.87119 | 0.58694 | 0.87203 | 0.59366 |
| **P-value** | 0.99371 | 0.93832 | | |

**Table S4.**

Significantly enriched GO terms (if any) for the nodes of the cluster dendrogram from Figure 4B. The top 2 terms (if any) for each node are presented in Figure 4B. The table below presents the complete list of terms for each node. Enriched terms were identified using the ClusterProfiler package for R, with an FDR cutoff of 0.05.

| Node | GO_Term | FDR |
|---|---|---|
| 1 | single-stranded DNA binding | 0.034275 |
| 1 | profilin binding | 0.038118 |
| 2 | cadherin binding | 0.006556 |
| 2 | RAGE receptor binding | 0.006556 |
| 2 | MHC protein complex binding | 0.009543 |
| 2 | S100 protein binding | 0.009543 |
| 2 | MHC class II protein complex binding | 0.013507 |
| 6 | antigen binding | 0.000000 |
| 6 | receptor ligand activity | 0.007538 |
| 6 | receptor regulator activity | 0.010464 |
| 6 | neuropeptide hormone activity | 0.010464 |
| 6 | cytokine activity | 0.010464 |
| 6 | cytokine receptor binding | 0.017554 |
| 6 | ammonium ion binding | 0.017554 |
| 8 | unfolded protein binding | 0.000070 |
| 8 | ATPase regulator activity | 0.000154 |
| 8 | heat shock protein binding | 0.000224 |
| 8 | chaperone binding | 0.000358 |
| 8 | ubiquitin protein ligase binding | 0.000358 |
| 8 | ubiquitin-like protein ligase binding | 0.000556 |
| 8 | DNA-binding transcription repressor activity, RNA polymerase II-specific | 0.001173 |
| 8 | cytokine binding | 0.003775 |
| 8 | protein N-terminus binding | 0.005781 |
| 8 | transcription corepressor activity | 0.010809 |
| 8 | platelet-derived growth factor receptor binding | 0.011551 |
| 8 | protein binding involved in protein folding | 0.011551 |
| 8 | nucleoside-triphosphatase regulator activity | 0.013652 |
| 8 | cytokine activity | 0.029022 |
| 8 | ATPase activator activity | 0.029022 |
| 8 | cytokine receptor activity | 0.029022 |
| 8 | ion channel binding | 0.029022 |
| 8 | enzyme activator activity | 0.029426 |

| Node | GO_Term | FDR |
|---|---|---|
| 8 | E-box binding | 0.031497 |
| 8 | chemokine binding | 0.031497 |
| 8 | protein heterodimerization activity | 0.036065 |
| 8 | ephrin receptor binding | 0.036628 |
| 9 | antigen binding | 0.000000 |
| 9 | MHC class II receptor activity | 0.010723 |
| 9 | MHC protein complex binding | 0.029652 |
| 9 | cytokine binding | 0.033592 |
| 9 | MHC class II protein complex binding | 0.033592 |
| 9 | cytokine activity | 0.043339 |
| 10 | cytokine activity | 0.020104 |
| 12 | tubulin binding | 0.041407 |
| 12 | signaling adaptor activity | 0.041599 |

**Table S5.**

List of surface markers analyzed. This list was generated by cross referencing the gene symbols and Cluster of Differentiation identifiers from the Cell Surface Protein Atlas (http://wlab.ethz.ch/cspa/) with the list of highly variable genes identified in our dataset.

| | | | | | |
|---|---|---|---|---|---|
| ABCB1 | CD27 | CD7 | FCGR3B | KIR3DL1 | THBD |
| ADAM8 | CD276 | CD72 | FCRL5 | KLRB1 | TLR4 |
| ATP1B3 | CD300A | CD74 | IFNGR1 | KLRC1 | TNFRSF17 |
| BTLA | CD300E | CD79A | IGF1R | LILRA1 | |
| CCR7 | CD36 | CD79B | IL17RA | LRP1 | |
| CD14 | CD37 | CD83 | IL18R1 | MRC1 | |
| CD163 | CD3D | CD8A | IL1R2 | MS4A1 | |
| CD164 | CD3E | CD8B | IL2RA | NCAM1 | |
| CD180 | CD3G | CD93 | IL2RB | NCR3 | |
| CD19 | CD4 | CD96 | IL2RG | NT5E | |
| CD1C | CD40LG | CD99 | IL4R | PDGFRA | |
| CD1E | CD44 | CR2 | IL6ST | PLAUR | |
| CD2 | CD48 | CSF2RB | IL7R | PTPRC | |
| CD200 | CD5 | CXCR1 | ITGA6 | SELL | |
| CD200R1 | CD52 | CXCR3 | ITGAL | SELPLG | |
| CD207 | CD53 | CXCR5 | ITGAM | SEMA4D | |
| CD22 | CD55 | DPP4 | ITGAX | SEMA7A | |
| CD24 | CD63 | FCER2 | ITGB1 | SIGLEC6 | |
| CD244 | CD68 | FCGR2A | ITGB2 | SIRPB1 | |
| CD247 | CD69 | FCGR3A | KIR2DL3 | SIRPG | |

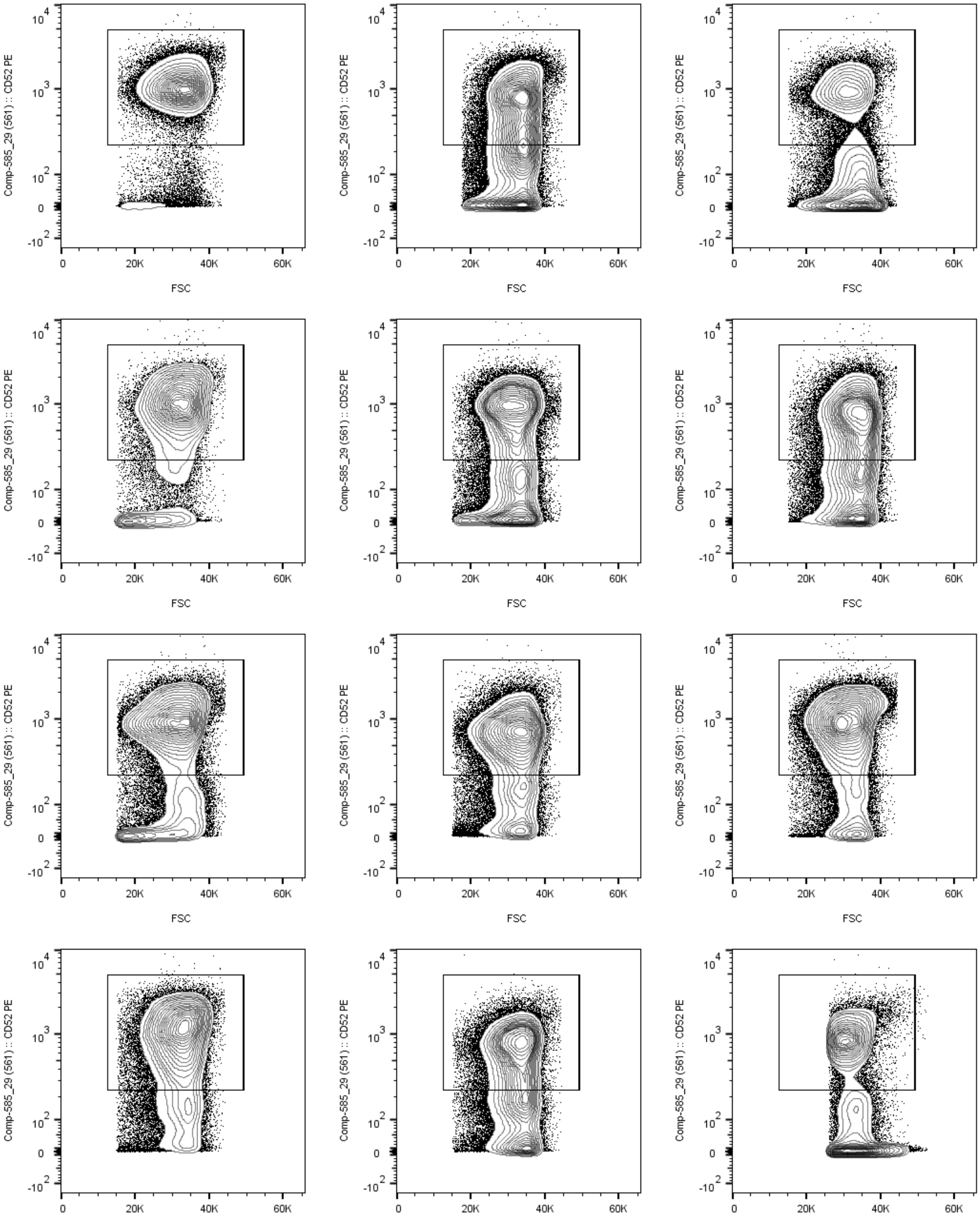

**Figure S1**: Representative scatter plots (CD52 vs. Forward Scatter) illustrating definition of the CD52+ gate (box). Cells shown are the CD3+/CD8+ NKT population. Twelve representative patients are shown.

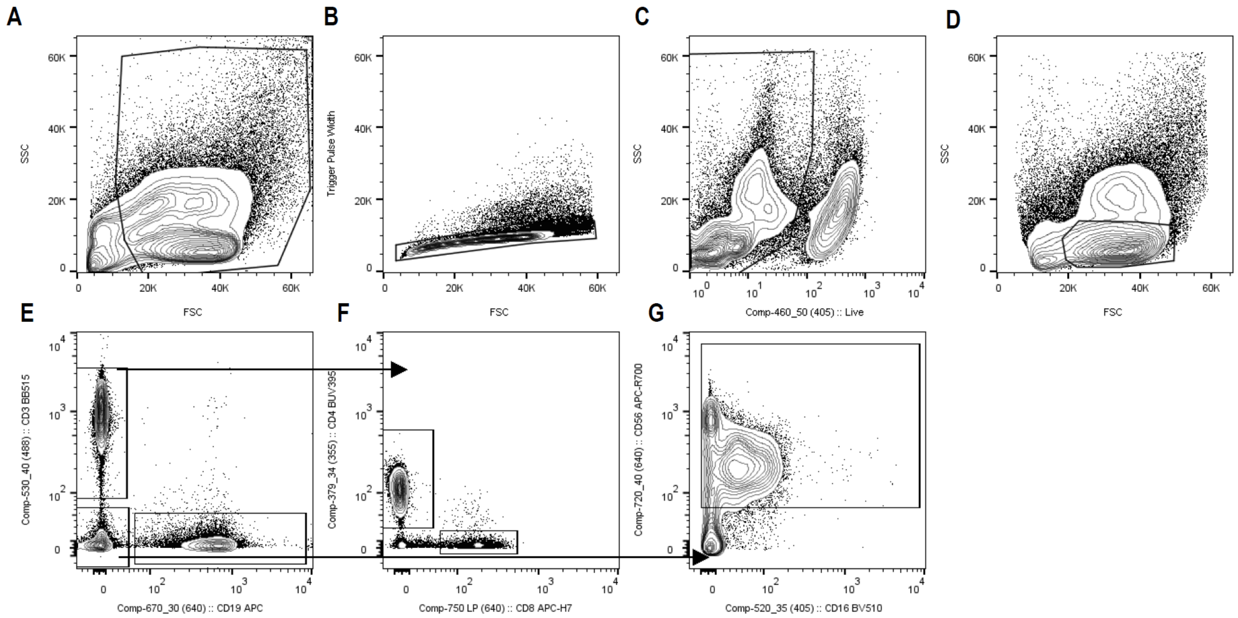

**Figure S2**: Representative scatter plots depicting the gating strategy used for FACS analysis of cells used to validate scRNASeq expression data (Fig. 3C). Non-debris was gated (A), followed by exclusion of potential doublets via FSC-w vs. FSC-h plot (B). From the previous gate, live cells were identified based on Fixable Viability Stain 450 fluorescence. From these live cells, a gate was drawn to select cells in the lymphocyte region based on FSC vs. SSC in the usual fashion (D). CD3+ (T cells) and CD19+ (B cells) populations were then easily distinguishable and gated (E). CD3+ cells were then divided into CD4+ and CD8+ T cell populations (F), while the CD3-/CD19- population was gated to identify CD56+ positive (NK) cells.

**Data S1. (separate file)**

Detailed description of the data analysis including annotated code to reproduce figures from the manuscript is provided in the file "final_analysis.html.pdf". Reproducing the analysis can be facilitated by downloading the github repository https://github.com/vanandelinstitute/va_ecls.

**Data S2. (separate file)**

The list of highly variable genes as defined by normalized dispersion used for the analysis shown in Figure 3B is provided in the file "Data_S2.tab"
